# Temporally robust occupancy frequency distributions in riverine metacommunities explained by local biodiversity regulation

**DOI:** 10.1101/2021.11.08.467721

**Authors:** Jacob D. O’Sullivan, J. Christopher D. Terry, Axel G. Rossberg

## Abstract

The mechanisms determining the distribution of the number of sites species occupy—the occupancy frequency distribution (OFD)—remain incompletely understood despite decades of research. Here we study OFD in fresh-water metacommunities in England. We show that OFD are largely time invariant despite continuous turnover in community composition and well described by log-series distributions. Both shape and temporal robustness are readily explained by a simple patch occupancy model with intrinsic regulation of local richness. The model is mathematically analogous to classical neutral theory, but concerns numbers of patches occupied instead of individuals present. The precise shape of the model’s log-series OFD is determined by a parameter representing the rate of colonisation from adjacent sites, which we estimate for the data. Our results support the view that meta-community structure reflects a dynamic steady state controlled by local ecological constraints. Interspecific trait differences underlie these constraints, but are unlikely to determine macroecological structure.

## 1 Introduction

Understanding the drivers of commonness and rarity in ecological communities, typically represented by the species abundance distribution (SAD), has been a central goal of community ecology for over seventy years (Preston, 1948). The determinants of commonness and rarity in *site occupancy*, represented by the occupancy frequency distribution (OFD), has motivated ecologists for even longer (Raunkiaer *et al.*, 1934). Site occupancy denotes the number of sites occupied by a given species, while the OFD describes the distribution of occupancy over a guild of species in a given landscape. Despite many decades of research, the mechanisms driving this fundamental ecological distribution remain incompletely understood.

In most assemblages the majority of species are observed in a minority of sites and the OFD is typically right-skewed (e.g. Tokeshi 1992; McGeoch & Gaston 2002; van Rensburg et al. 2000; Bossuyt et al. 2004; Heatherly et al. 2007; Astorga et al. 2014; Heino 2015; Boieiro et al. 2018; Renner et al. 2020). Bimodal OFDs, with modes at both the highest and lowest occupancy, have also been reported (e.g. Storch & Šizling 2002), but there are indications that such bimodality may reflect sampling resolution rather than the particularities of community assembly (van Rensburg *et al.*, 2000; Bossuyt *et al.*, 2004; Heatherly *et al.*, 2007). Here we focus on the spatial and taxonomic sampling scales for which unimodality is understood to dominate.

A still-influential review of OFD research by McGeoch & Gaston (2002) identified a number of biological mechanisms plausibly responsible for the shape of the distribution: interspecific variation in niche breadth, interspecific variation in dispersal ability, and site occupancy dynamics described in Hanski’s (1982) classical joint metapopulation model. Alternative drivers were considered but relate more to sampling regime than to underlying biology. We argue that these explanations, while widely accepted, cannot sufficiently explain the shape of the OFD.

Environmental niche widths are likely to influence occupancy at the species scale, with habitat specialists and generalists typically of low and high occupancy respectively (Brown, 1984; Brown *et al.*, 1995; Denelle *et al.*, 2020). A positive relationship between niche breadth and *range size—*generally found to correlate with occupancy—has been reported across taxonomic groups and spatial scales (Slatyer *et al.*, 2013). However, niche width and range size estimates are frequently made by inspection of the same data (Carscadden *et al.*, 2020) and as such the association may often be tautological.

Moreover, the apparent relationship between niche breadth and site occupancy alone does not explain the *shape* of the OFD. This would also require that narrow niche widths are substantially over-represented in nature. Evidence for such a tendency is weak. Pooling data at the global scale, Belmaker *et al.* (2012) find a numerical dominance of diet and habitat specialists in birds. However, a positive association between latitude and niche breadth (Stevens, 1989; Pagel *et al.*, 1991; Ruggiero, 1994) contrasts with the negative latitudinal trend in species richness (Pianka, 1966). Together these mean that the bias toward narrow niche widths is likely exaggerated at the global scale compared to the narrower latitudinal ranges at which OFD are reported. At regional scales, weak right-skew in environmental tolerance has been reported in riverine macroinvertebrates (Mykrä & Heino, 2017). However, this contrasts with a symmetric distribution in freshwater macrophyte niche breadths (Kennedy *et al.*, 2017) and an over-representation of generalists in a regional-scale plant assemblage (Boulangeat *et al.*, 2012). Thus, clear and consistent evidence of right-skew in the distribution of niche widths explaining the ubiquitous right-skewed OFD has not been found.

Alternatively, high occupancy could plausibly be associated with good dispersal ability (Collins & Glenn, 1997). While evidence for a mechanistic link between dispersal ability and total range size is inconclusive (Sheard *et al.*, 2020), positive relationships have been shown (Capurucho *et al.* 2020; Böhning-Gaese *et al.* 2006; Luo *et al.* 2019; Laube *et al.* 2013; Sheard *et al.* 2020 but see Lester *et al.* 2007). Right-skew, sometimes strong, has been found in the distribution of proxies for dispersal ability or estimated dispersal rates (Schloss *et al.*, 2012; Tamme *et al.*, 2014; Sarremejane *et al.*, 2020; Sheard *et al.*, 2020). Again, however, differences in typical dispersal abilities between biomes (Chen *et al.*, 2019; Álvarez-Noriega *et al.*, 2020; McCaslin *et al.*, 2020) mean that global trait databases are likely to miss-represent landscape-scale distributions. Moreover, OFD are generally reported for guilds of species for which common proxies for dispersal ability based on body- or propagule-size do not vary substantially. A correlation between such proxies and *within* guild occupancy has, to our knowledge, not been shown.

A uniquely phenotypic explanation for the shape of the OFD would imply that species tend to occupy roughly consistent numbers of sites through time in the absence of strong environmental change. In fact, species’ site occupancy in contemporary England is extremely variable in time (Wayman *et al.* 2022, and below). Hanski (1982) proposed a family of metapopulation models with fluctuating parameters that lead to fluctuations in site occupancy and distributions similar to those observed. As an explanation, however, Hanski’s (1982) theory is incomplete since it does not specify the source of fluctuations or explain why fluctuations have the magnitude required to reproduce observations. Hanski (1982) considered that competition between species might moderate model parameters, but the process driving the model appears to arise at population-level.

Here we test an alternative hypothesis, that the shape of the OFD is a fundamental outcome of an ecosystem subject to intrinsic constraints on biodiversity. This *ecological* explanation does not depend on interspecific heterogeneities, but instead on strong constraints on site-level species richness reported in both theoretical (Drossel *et al.*, 2001; Yoshida, 2003; Pawar, 2009; Rossberg, 2013; O’Sullivan *et al.*, 2019) and empirical studies (Brown *et al.*, 2000; Parody *et al.*, 2001; Magurran & Henderson, 2010; Magurran *et al.*, 2015; Dornelas *et al.*, 2014; Gotelli *et al.*, 2017; Vellend *et al.*, 2017). Such constraints occur at the local scale, the outcome of limits on the structure and diversity of local ecological assemblages. Evidence of local processes driving metacommunity patterns in biodiversity would represent an important step toward the elusive theoretical union of local and regional community scales (Rabosky & Hurlbert, 2015; Harmon & Harrison, 2015).

A key signature of such an ecological mechanism would be the detection of *temporally robust empirical OFDs despite continuous change in the identities and occupancies of the constituent taxa*. We test for this signature here. We find that the shape of the OFD for riverine communities is indeed typically time invariant despite pervasive temporal turnover. We interpret the empirical OFDs using a simple patch occupancy model which includes intrinsic constraints but in which colonisation and extinction rates are independent of species identity. The steady state OFD of this simple model is a log-series with a single fitting parameter which summarises the internal colonisation rate for the metacommunity. We find a consistently good fit of the parameterised log-series, which outperforms a variety of plausible alternative distributions, to the empirical OFD. We conclude that the simple model we propose may offer substantial opportunities for inferring the mechanisms determining macroecological structure, including patterns in spatial and temporal occupancy.

## 2 Methods

### 2.1 Compiling the database of temporal OFDs

We studied occupancy patterns in macroinvertebrate (I), macrophyte (P) and diatom (D) surveys compiled by the Environment Agency (EA) of England^1^. The database covers the whole of England at good spatial resolution (Fig. 1) and includes up to 30 years of macroinvertebrate observations (somewhat less for the other two taxonomic groups). We retained only observations identified to species or genus level. Of these, taxa identified to genus level made up 31%, 30%, and 11% of the I, P and D datasets. Detailed descriptions of the sampling procedures can be found in EA documentation^2 3 4^.

**Figure 1:**
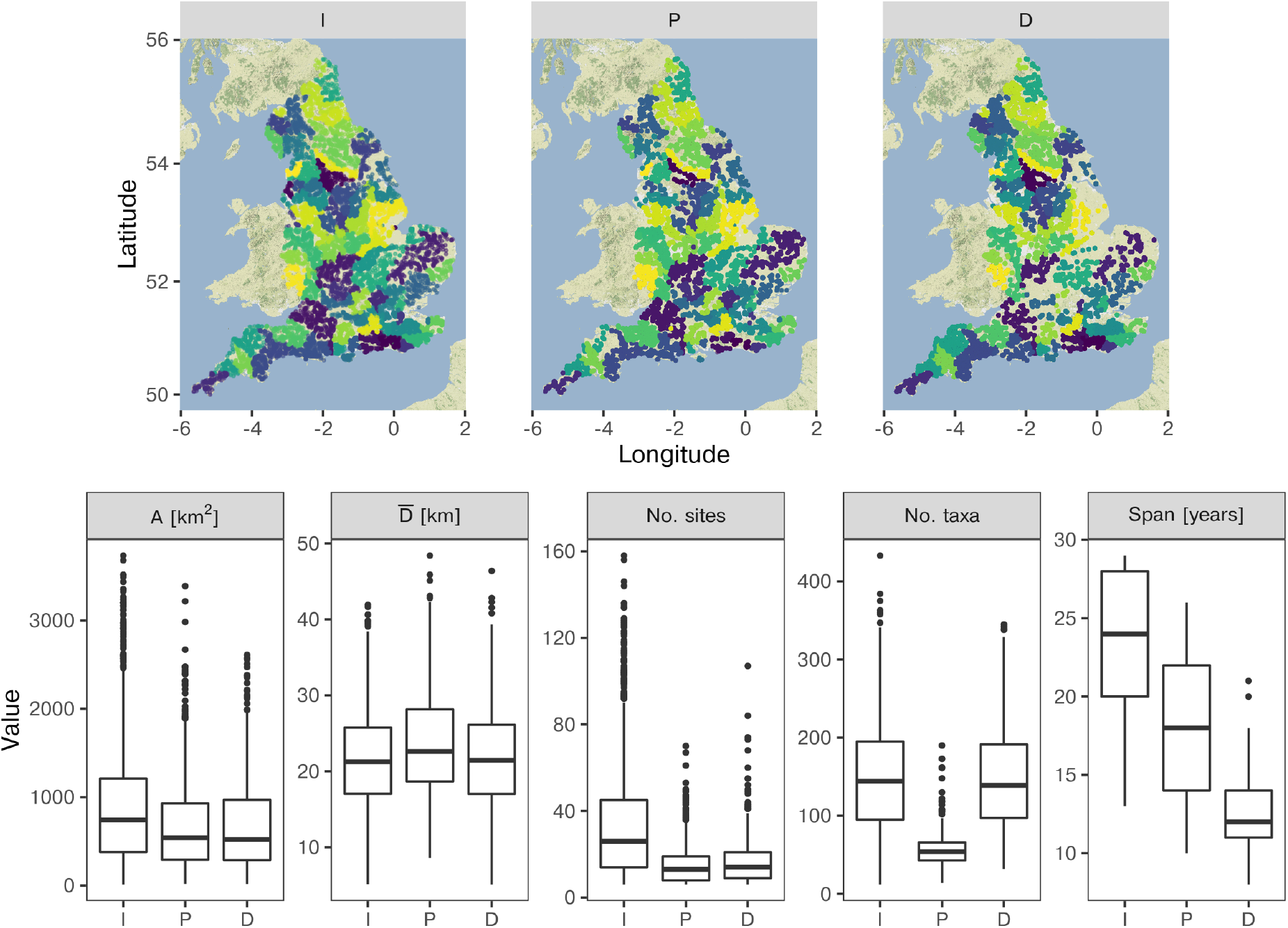
Descriptive statistics for the EA database following filtering and grouped by Management Catchment. Top row: The locations of sites in the macroinvertebrate (I), macrophyte (P) and diatom (D) surveys. Points correspond to sites, colours to management catchment. Bottom row: The area of the convex hull bounding sites, mean inter-site distance, number of sites, the number of taxa observed in individual catchment years, and the temporal span of the catchment time series, each plotted in the standard Tukey representation (Tukey, 1977). In all cases, figures represent the datasets after filtering.

The EA divides riverine ecosystems in England into 10 River Basin Districts, each of which contains between 2 and 20 Management Catchments. Management Catchment allocation was extracted from the EA Catchment data explorer^5^ and this information was merged with the database of observations. Management Catchments are delineated by the EA based on analysis of flow and terrain data as well as hydrological modelling, “ensuring that [Water Framework Directive] rivers do not intersect boundaries”^6^. Here, we study each Management Catchment as a separate metacommunity. The EA also define an additional higher-resolution organisational scale referred to as Operational Catchments (1 - 10 per Management Catchment, median 3). Repeating the key analyses described below for Operational Catchments, we found the conclusions to be unimpacted.

We mitigate potential impacts of insufficient sampling effort by applying a suite of filters to the data (see Supplementary information Sec. S1, Fig. S1). Following filtering, we retain 83, 52 and 48 catchments (I, P and D respectively), sampled over a period of up to 30 years, resulting in 1804, 731 and 588 unique OFDs. The location of each site and a variety of relevant catchment-scale properties are shown in Fig. 1.

### 2.2 Testing for bimodality in the OFDs

Our method for describing the shape of the OFD assumes unimodality—specifically, a single mode in the OFD at low occupancies—and broad consistency in the qualitative shape of the OFD across space and time, such that quantitative change is captured by the OFD’s lowest statistical moments. If this were not the case, a more *ad hoc* analysis would be required. We therefore assessed the overall shape of the OFDs for each catchment-year by testing for bimodality using Tokeshi’s (1992) method, asking whether the observed numbers of species in both the left- and right-most 10% of the occupancy classes each separately exceed null expectation for a uniform ODF at 95% confidence level.

Significant bimodality—greater than expected richness at both extremes of the OFD based on the assumption of uniformity—was detected in 4%, 15% and 26% of the catchment-years for the I, P and D datasets prior to filtering. The remaining OFD all had single modes in the occupancy class representing up to 10% of survey. After filtering the percentages dropped to 0.2% 0.1% and 6% respectively (Fig. S2). This suggests that, for these data at least, bimodality in the OFD is predominantly a result of limited sampling rather than a biological signal. Considering this broad consistency in the qualitative shape of the OFDs, we restrict our analysis to the first three moments of these distributions to compare them between catchments-years.

### 2.3 Catchment-scale turnover in composition and occupancy

The *absence* of change in OFD over time is a phenomenon requiring explanation only in the case that it occurs despite meaningful ecological change over the time series. We investigated the temporal trends in community turnover using two measures of compositional similarity. The first was the average change in Jaccard similarity (Jaccard, 1912) in *catchment* composition over time (i.e. considering all sites in the catchment-year as a single sample). The second was the average change in Bray-Curtis similarity (Bray & Curtis, 1957) in the vectors describing the proportional site occupancy for each species over time. In both cases these were studied as functions of time lag irrespective of directionality (Fig. S3).

The EA biomonitoring program aims to assess the ecological status of a waterbody or set of waterbodies, according to a variety of qualitative and quantitative indices. It was not designed with metacommunity theoretical analyses such as the present one in mind. Particularly problematic, the sites selected for monitoring can vary substantially between years for many catchments. Catchment-scale measures of turnover must, therefore, be interpreted with caution. For comparison, we also studied the decay in Jaccard similarity for those local sites that were sampled a minimum of 5 times (230, 92 and 113 sites for I, P and D datasets respectively), limiting our inspection to a maximum lag of 10 years to account for likely changes in methodology and/or personnel over the period. Additionally, in supplementary Sec. S6 we mathematically consider the question to what extent apparent turnover can result from the selection of different sampling sites. This analysis shows that, assuming the theoretical OFD proposed below, there is an upper limit to such apparent turnover in the case that no actual biological change occurs at the catchment-level. We compared the observed turnover to this theoretical limit to further test whether observed change can be explained by the sampling protocol employed.

### 2.4 Testing for temporal trends in empirical OFDs

For analyses of temporal trends in the OFD we focus on the *proportional* site occupancy to account for the fact that number of sampling sites was not fixed through time for a given catchment. We tested whether the shape of the OFDs varied over time by analysing temporal fluctuations in the mean, variance and skewness of the distribution for each catchment. To account for intrinsic dependencies between subsequent data in time series, the existence of a trend in each of these descriptive statistics was tested using the Mann-Kendall test (Mann, 1945; Kendall, 1948b) and the slope of the trend assessed using Sen’s estimator (Theil, 1950; Sen, 1968). We also tested for stationarity in the time series using Augmented Dickey Fuller (ADF) (Fuller, 2009) and KPSS tests (Kwiatkowski *et al.*, 1992). The ADF fits an autoregressive model of arbitrary order to a time series and tests for the presence of a unit root, indicative of non-stationarity. In the ADF, the presence of a unit root (non-stationarity) is the null hypothesis. The KPSS is analogous, however the null hypothesis is the absence of a unit root (stationarity).

### 2.5 A locally saturated patch occupancy model

We model the OFD at the catchment scale using the Locally Saturated Patch Occupancy Model (LSPOM) defined in Box 1. The LSPOM incorporates into a conventional patch-occupancy formalism (Levins, 1969; Day & Possingham, 1995) a robust constraint on local species richness. This constraint is understood to be the outcome of ecological structural instability, which arises with increasing species richness in complex ecological interaction networks (Rossberg, 2013). In the LSPOM, for simplicity and analytic tractability, these complex networks are entirely implicit.

##### Box 1: The Locally Saturated Patch Occupancy Model and its steady state

The Locally Saturated Patch Occupancy Model (LSPOM) builds on the patch-occupancy paradigm (Levins, 1969), while also describing stochasticity arising from the discreteness of colonisation and extirpation events (Day & Possingham, 1995). The LSPOM makes two additional assumptions: each patch *i* of a metacommunity, extending over *N* patches, can sustain a maximum species richness *α_i_* and all patches are saturated with species, i.e. reach this limit *α_i_*. As a result, the metacommunity hosts 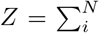 populations. Time in the model is measured by the number of successful invasions of new species into the metacommunity. New invasions lead to replacement of one of these *Z* populations, chosen at random, by a population of the invading species. During each time step, any of the *Z* resident populations colonises another patch with a probability *m/Z*, where *m* is called the ‘mixing rate’. The colonised patch i is chosen at random with probability *α_i_/Z*. Each such colonisation leads to extirpation of a resident population from the colonised patch. Thus, on average m colonisations from within the metacommunity occur in each time step and *m* + 1 extirpations.

Simulations (see Supplementary information Sec. S2) confirm that in the model steady state most species occupy 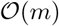 patches. For an analytic treatment assuming that *N* is large compared to typical occupancy and *Z* ≫ max_*i*_ *α_i_*, one can therefore disregard the constraint that each species has at most one population in each patch and instead simply keep track of the number of populations formed by each species in the model. The mean OFD in the model steady state is then obtained as the log-series distribution

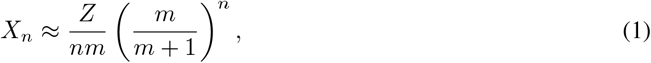

implying an average species occupies of *m*/ ln(1 + *m*) patches. For a detailed elementary derivation, see Supplementary Information Sec. S3. In Fig. S4 we demonstrate how the mixing rate parameter m in direct simulations of the LSPOM can be recovered by fitting the steady-state OFD to Eq. (1).

The steady state of the LSPOM is a (unimodal, right-skewed) log-series OFD (Box 1, supplementary Sec. S3). The shape of the log-series OFD is controlled by a single parameter, which we term the ‘mixing rate’ *m.* The parameter can be understood as the internal colonisation rate—the rate at which local populations colonise additional sites within the metacommunity—in units of the rate of successful invasions into the metacommunity. Since the steady-state OFD of the LSPOM is controlled by *m* alone, it can readily be fitted to observed OFDs (Box 1 and Supplementary Secs. S4-S5). We computed, for each catchment and taxonomic group, the empirical OFD averaged over the filtered time series. We then obtained simple least-square fits of the theoretical log-series OFD, denoting the best-fitting mixing-rate parameter *m* by *m*_raw_.

### 2.6 Correcting for incomplete observation

The estimated log-series parameter, *m*_raw_, must be corrected for false-negatives (unobserved species). In Supplementary Information Sec. S4 (see also Fig. S5) we describe the procedure for estimating *ω*, the detection probability and show that observation of species with a fixed false-negative rate 1 – *ω* transforms the theoretical OFD in Box 1 into an OFD with the same functional form but with *m*_raw_ replaced by *mω*. We therefore computed for each catchment and taxonomic group a mixing rate corrected for false negatives as

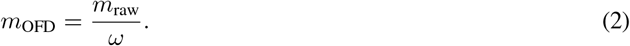

Similarly, the mixing rate estimates should be additionally corrected to account for the case that a proportion *q* < 1 of the sites of the ‘actual’ metacommunity is surveyed. Again, the predicted functional form of the OFD remains unchanged, but with a linear correction by *q* to *m*_OFD_. Since we have no clear route to estimating *q* from the available data, we assume here that *q* ≈ 1. Further work involving rigorous quantitative assessment of the estimated mixing rate should involve estimation of *q*.

### 2.7 Comparative assessment of the log-series OFD

Quantitative validation of macroecological theory is a major challenge (McGill, 2003). Here we test the LSPOM log-series (hereafter log-series) OFD after averaging the richness observed in each occupancy class over the time series. This averaging procedure reduces noise due to observation error (rather than actual fluctuations in occupancy) and is justified in light of the comparatively long time series (Fig. 1). We compared the time-averaged OFDs to the log-series using Pearson’s (1900) chi-square goodness-of-fit (GOF) analysis, binning the distribution according to three different schemes: 10 or 5 arithmetically spaced, or 10 logarithmically spaced bins. For comparison, we also fit four ‘purely statistical’ distributions previously employed to explain empirical SAD (McGill *et al.*, 2007) and repeated the GOF analysis. These were the log-normal, gamma, negative binomial and geometric distributions with parameters estimated by maximum likelihood using the R function **fitdist** (Delignette-Muller & Dutang, 2015). At the national scale we compared the catchment-time-averaged OFD for each taxonomic group independently to the same theoretical distributions. In this case, chi-square testing is hard to justify. Instead, we compared the maximum absolute difference between the average observed and predicted richness in occupancy classes.

### 2.8 Further investigation of LSPOM estimates: a demonstration of concept

Given that strongly right-skewed distributions generated by a variety of mechanisms are difficult to empirically distinguish, it is crucial to develop theories that move beyond simple curve fitting and test for additional patterns predicted by a specific explanatory model (McGill, 2003). Here we propose such a test, based on quantitative analysis of catchment-scale turnover rates. Unfortunately, due to the methodological issues described in Sec. 2.1, turnover rates cannot be determined with sufficient confidence to perform this test. As a proof of principle, however, we demonstrate the required calculations in Sec. S7.

## 3 Results/Discussion

### 3.1 Temporal trends in catchment OFDs

Both the Jaccard similarity in catchment composition and Bray-Curtis similarity of proportional occupancy displayed clear signs of temporal turnover. In Figs. 2A and S7 we show the decay in these two similarity measures. At the local site scale the tendency for Jaccard similarity to decay with temporal lag was also pervasive and remarkably consistent across sites (Fig. S6). Furthermore, we found that, for 93, 76 and 84% of the catchments for I, P and D datasets respectively, the observed compositional change exceeded the theoretical maximum predicted in the absence of true turnover (see Supplementary information Sec. S6). Thus, we conclude that catchment-scale turnover is indeed a general property of riverine communities in England, rather than a methodological artefact.

**Figure 2:**
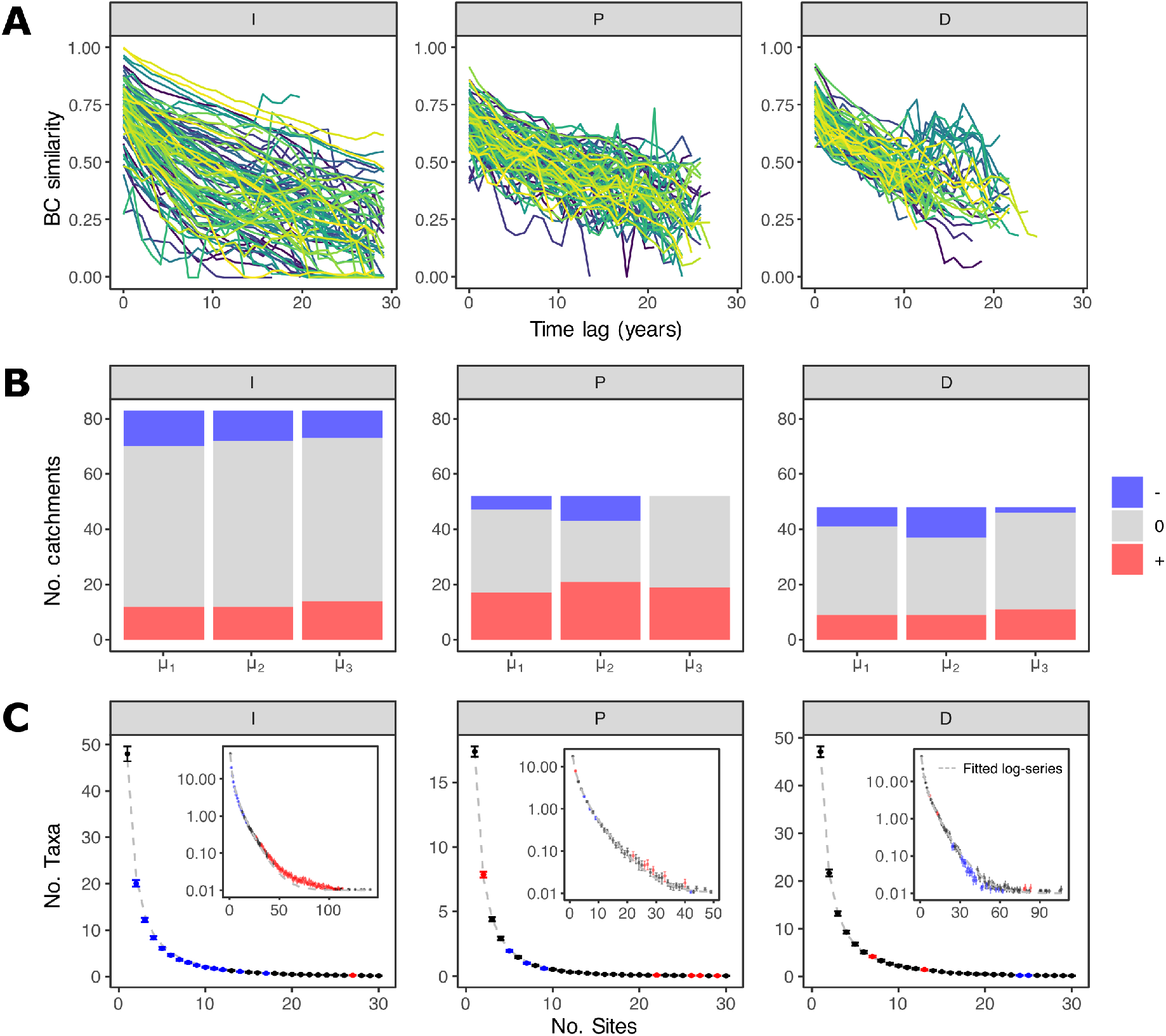
Compositional turnover, temporal robustness in the OFD, and fitting the log-series to the national averages. **A**: Turnover in composition and occupancy at the metacommunity scale shown as the average BC similarity of the proportional site occupancy vectors plotted against time lag. Local (Fig. S6) and metacommunityscale turnover in composition (Fig. S7) and occupancy was pervasive in each of the datasets, and greater than predicted due to site variation alone for the majority of catchments. **B**: The outcome of time series analysis of the first three moments of catchment OFDs. For each of the mean (*μ*_1_), variance (*μ*_2_) and skewness (*μ*_3_), in all three taxonomic groups, the majority of catchments showed no significant temporal trend (95% confidence level), with the single exception of the variance in macrophyte OFDs. Colour represents the sign of the Sen’s trend, with grey indicating no detectable trend by Mann-Kendall test. **C**: OFDs obtained by averaging across catchments and years, fitted by the log-series. Error bars represent standard error of the means. Blue (red) points indicate occupancy classes of richness less (greater) than log-series model prediction. For visual clarity the OFDs are truncated to 30 sites. Insets show the full range of observed occupancy classes on a logarithmic richness scale, after adding 10^-2^ to avoid singularities. The estimated mixing rates for these composite OFDs (without false-negative correction) were 11.96, 7.03 and 10.67 for the I, P and D datasets, respectively. Alternative model fits to the national-average OFD are shown in Fig. S15.

Despite this pervasive tendency for catchment composition to turn over through time, we detected significant temporal trends in mean, variance or skewness of the catchment OFDs for only a small proportion of the catchments. In Figs. S8-S10 we show the linear trends (Sen’s slope) of the first three moments over time for 12 random catchments, and in Fig. 2B the number of catchments for which positive, negative or no trend were detected. For macroinvertebrates and diatoms in particular, a substantial majority of catchments show no temporal trend in the first three moments of the OFD. The only exception to the numerical dominance of catchments with no detectable temporal trend in shape was in the variance of macrophyte OFDs. The proportion of significant trends overall is higher than would be expected by random chance, however these cases are exceptional nonetheless when contrasted with the evidence for metacommunity turnover. It is conceivable that the catchments for which significant trends were detected are responding to multidecadal environmental change of natural or anthropogenic origin.

Applying the same analyses to the data clustered in Operational Catchments, we found again found evidence of pervasive turnover. Unsurprisingly, data at the smaller organisational scale tend to be more noisy. This is due the fact that sample sizes (number of sites, number of taxa, number of catchment-year retained after applying the same filtering criteria) were considerably smaller for Operational catchments. Again, however, a substantial proportion of Operational Catchments (60-71% for I and D, 50-66% for P dataset) show no detectable trend in any of the three moments of the empirical OFD.

We then tested for stationarity in the time series of the moments (Fig. S11). For the majority of catchments and in all three datasets, using ADF we did not find evidence of stationarity in the time series of the mean, variance and skewness of the catchment ODFs (95% confidence level). However, using the KPSS test, we also did not find evidence of non-stationarity. Across catchments, evidence for stationarity tended to be stronger for variance than for mean and skewness (Fig. S11). Thus, we cannot conclusively confirm that fluctuations in the shape of the OFD can be modelled as stationary processes.

Our tests for stationarity in the first three moments of the catchment OFDs, based on standard methods, were inconclusive: we find temporally ‘robust’ OFD but cannot demonstrate or disprove stationarity. Stationarity tests have previously been used to evidence intrinsic regulation of species richness (Gotelli *et al.*, 2017). There is good reason to assume that metacommunity dynamics occur at a rate slower than local community dynamics (O’Sullivan *et al.*, 2019), meaning substantially longer time series would be required to conclusively demonstrate stationarity in metacommunity properties. In fact, the decline of compositional similarity with increasing time lag shown in Figs. 2A and S7 suggests that the 15 - 30 year period for which data are available corresponds in many cases to around one ‘half-life’ of catchment-scale turnover (similarity drops to ~ 0.5). Even multi-decadal time series might therefore be too short for robust inference regarding stationarity of fluctuations. Since the OFD is a *distribution* rather than a scalar property like species richness, we can arguably rely on the similarity of this distribution across distant catchments and taxonomic groups as *prima facie* evidence of intrinsic regulation via some common mechanisms.

### 3.2 Patch-occupancy modelling of the OFD

In view of the tendency for OFD to be robust in time, we fit their temporal averages to the theoretically predicted log-series as well as four alternative distributions. In Figs. S12-S14 we show the time-averaged OFD fit by the log-series for each taxonomic group in 25 random catchments. The shapes of OFD are highly consistent across catchments and taxonomic groups. The number of catchments and percent of the available data for which the time-averaged OFD could plausibly be described as samples from the five theoretical distributions are shown in Table 1 (Pearson’s χ^2^ GOF analysis, *p* > 0.05). Of the catchments retained after filtering, 75-100%, depending on taxonomic group and binning, match the log-series prediction. The log-normal distribution also matches a relatively large proportion of the available data (14-100%), while the gamma distribution frequently captured the shape for the macrophytes. The geometric and negative binomial distributions matched a substantially smaller proportion of the empirical OFD (0-55.8%).

**Table 1:** The number (percentage) of catchments for which the empirical OFD matched theoretical predictions (Pearson’s χ^2^, *p* > 0.05). OFD were grouped in 10 (χ^2^ *A*) of 5 arithmetically (χ^2^ *B*), or 10 logarithmically spaced bins (χ^2^ *C*). We tested the LSPOM log-series (LS), as well as log-normal (LN), gamma (Ga), geometric (Geo) and negative binomial (NB) distributions (table ordered by row sums).

| | $\chi^2 A$ | | | $\chi^2 B$ | | | $\chi^2 C$ | | |
| --- | --- | --- | --- | --- | --- | --- | --- | --- | --- |
|  | I | P | D | I | P | D | I | P | D |
| LS | 83 (100%) | 51 (98.1%) | 41 (85.4%) | 82 (98.8%) | 50 (96.2%) | 36 (75.0%) | 82 (98.8%) | 51 (98.1%) | 40 (83.3%) |
| LN | 78 (94.0%) | 52 (100.0%) | 36 (75.0%) | 53 (63.9%) | 52 (100.0%) | 23 (47.9%) | 25 (30.1%) | 52 (100.0%) | 7 (14.6%) |
| Ga | 13 (15.6%) | 47 (90.4%) | 6 (12.5%) | 9 (10.8%) | 47 (90.4%) | 4 (8.3%) | 1 (1.2%) | 40 (76.9%) | 0 (0.0%) |
| Geo | 3 (3.6%) | 29 (55.8%) | 0 (0.0%) | 0 (0.0%) | 8 (15.4%) | 0 (0.0%) | 0 (0%) | 16 (30.8%) | 0 (0.0%) |
| NB | 1 (1.2%) | 20 (38.5%) | 1 (2.1%) | 1 (1.2%) | 17 (32.7%) | 0 (0.0%) | 0 (0.0%) | 4 (7.7%) | 0 (0.0%) |

Averaging the time-averaged OFD across catchments we also find a strikingly good fit to the log-series. In Fig. 2C blue (red) points indicate occupancy classes for which the the log-series prediction (grey line) is greater (less) than the range indicated by the standard error of the mean (SEM, indicated by error bars) computed with catchments as samples. We find a weak tendency for the richness of low occupancy classes to be overestimated (blue points), and intermediate-to-high classes to be underestimated for the macroinvertebrates (red points) by the model. This suggests log-series OFD may not account for infrequent, extremely high occupancy taxa observed in the macroinvertebrate dataset. This could reflect biology absent from the LSPOM, for example the signature of extremely mobile or competitively dominant taxa. The maximum absolute difference between the empirical and theoretical OFDs at the national scale are shown in Table 2. For the log-series, deviations between observed and expected richness in occupancy classes ranged from 0.49-2.88 species, compared to 2.26-15.26 for the next best fitting distribution, the log-normal. For completeness we plot the catchment-time-averaged OFDs with the best fitting contender models in Fig. S15.

**Table 2:**
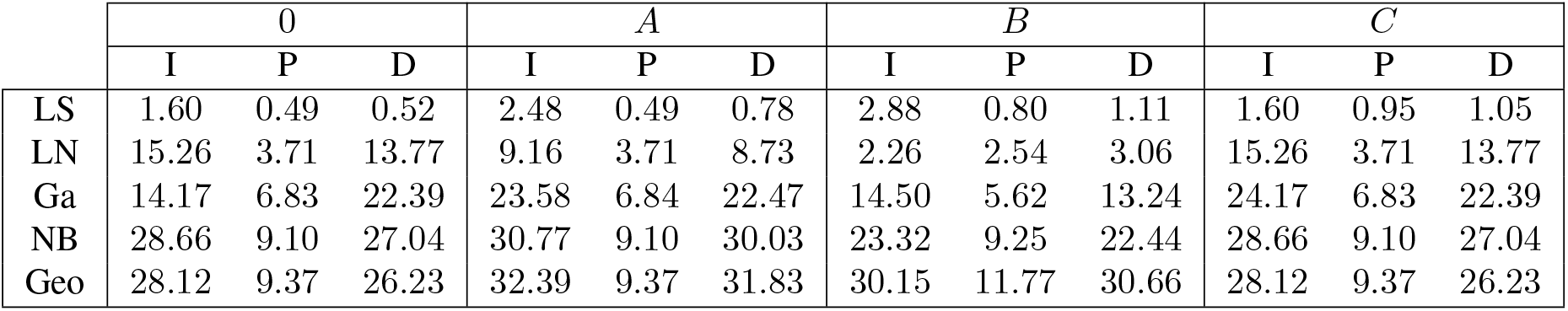
The maximum absolute difference (number of species) between observed and predicted catchment-time-averaged OFD based on the various theoretical distributions (ordered by row sums, note the difference in ordering relative to Table 1). OFD were binned in integer numbers of sites (0) and, for comparison, according to the binning scheme employed previously (*A-C*).

|  | 0 |  |  | *A* |  |  | *B* |  |  | *C* |  |  |
| --- | --- | --- | --- | --- | --- | --- | --- | --- | --- | --- | --- | --- |
|  | I | P | D | I | P | D | I | P | D | I | P | D |
| LS | 1.60 | 0.49 | 0.52 | 2.48 | 0.49 | 0.78 | 2.88 | 0.80 | 1.11 | 1.60 | 0.95 | 1.05 |
| LN | 15.26 | 3.71 | 13.77 | 9.16 | 3.71 | 8.73 | 2.26 | 2.54 | 3.06 | 15.26 | 3.71 | 13.77 |
| Ga | 14.17 | 6.83 | 22.39 | 23.58 | 6.84 | 22.47 | 14.50 | 5.62 | 13.24 | 24.17 | 6.83 | 22.39 |
| NB | 28.66 | 9.10 | 27.04 | 30.77 | 9.10 | 30.03 | 23.32 | 9.25 | 22.44 | 28.66 | 9.10 | 27.04 |
| Geo | 28.12 | 9.37 | 26.23 | 32.39 | 9.37 | 31.83 | 30.15 | 11.77 | 30.66 | 28.12 | 9.37 | 26.23 |

The estimated log-series parameter, *m*_OFD_, without accounting for spatial coverage (unknown), ranged from 4.06-48.67 (median 17.63), 5.64-38.77 (median 10.55) and 7.36-77.45 (median 20.92) for the I, P and D datasets respectively (Fig. 3). The estimated mixing rates for the catchment-time-averaged OFD were 11.96, 7.03 and 10.67 for the I, P and D datasets respectively, broadly consistent with the median estimates at local catchment scale. Comparative analysis suggests at a minimum that macrophytes tend to colonise adjacent sites in riverine metacommunities at a lower rate than macroinvertebrates or diatoms, an inference that could plausibly be validated using alternative methods, e.g. by comparing demographic rates between taxonomic groups. In Supplementary Sec. S7 we show how these values could be compared to alternative estimates based on temporal turnover dynamics. We find that these estimates differ by a factor of between 4 and 8 on average, a difference we might plausibly expect given the simplifying assumption that *q* =1, and the impact of inter-annual site variation on estimated catchment turnover rates.

**Figure 3:**
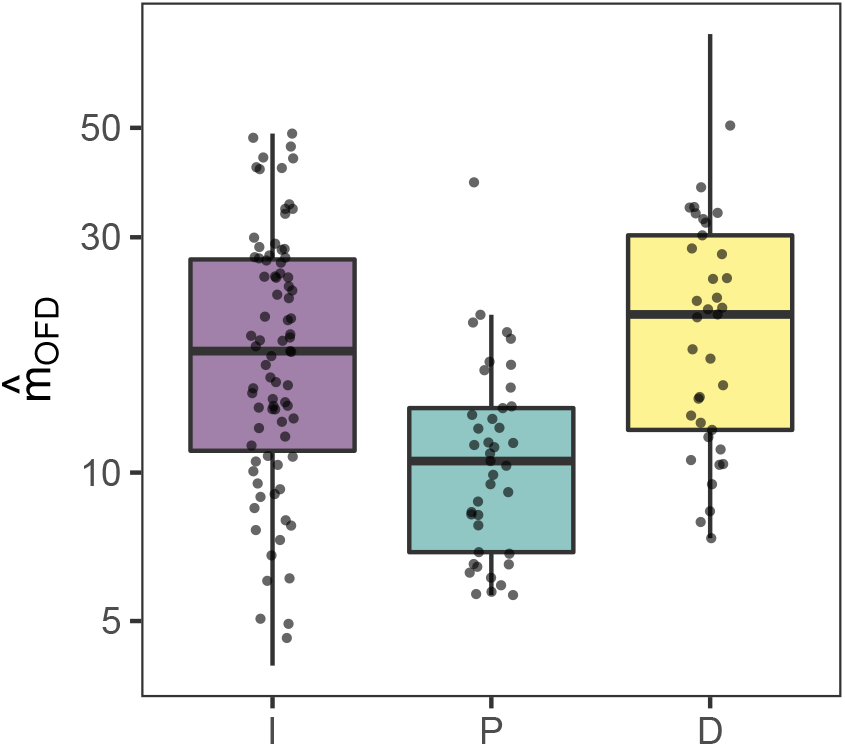
Log-series parameter *m*_OFD_ estimated for the time-averaged catchment OFDs. The mixing rate *m*_OFD_ describes the internal rate of site colonisations in the empirical metacommunities. Here we report the estimated values for the three datasets after correcting for false-negatives. Work to quantitatively validate these parameter estimates is ongoing, with an example procedure described in the Supplementary Sec. S7.

McGill 2003 argued that simple curve matching is a weak test of the validity of theory in macroecology for various reasons: Multiple theories may predict the same distribution (Pielou, 1977). To date, we know of no alternative theory which has predicted a log-series OFD. Theories with many free parameters are extremely difficult to disprove. The LSPOM includes a single parameter which has a clear (if composite) biological meaning. Different goodness of fit tests can favour different explanatory models for the same data. Here we compute GOF statistics for the log-series but caution that these alone should not be interpreted as sufficient evidence of the plausibility of the model. We note, however, that compared to comparable tests for SAD (McGill, 2003) a substantially higher percentage of replicates (catchments) conform to the log-series expectation of the OFD. Finally, while currently inconclusive, we have developed a plausible method for validating the log-series according to the additional criterion proposed by McGill 2003: that theory in macroecology should quantitatively predict multiple ecological phenomena simultaneously.

## 4 Conclusions

Despite decades of research, the mechanisms responsible for the shape of the OFD have remained poorly understood. Researchers often lean on incomplete explanations targeted at the species level—niche width or dispersal heterogeneities—but ignore the fact that on their own such heterogeneity sheds little light on the cause of the typically right-skewed shape of observed OFDs. With the present study we provided compelling evidence in favour of a fundamental macroecological mechanism underlying this pattern. We have successfully modelled observed OFD as the outcome of a stochastic colonisation-extinction process with equal probabilities for all species—i.e. independent of the phenotypic differences commonly promoted in the OFD literature—assuming locally saturated assemblages. This implies that interspecific differences are unlikely to play a dominant role in determining this fundamental pattern. Our proposed model improves on previous attempts to distil the basic biology structuring macroecological assemblages in its simplicity and the readiness with which it can be fitted to empirical data. The LSPOM implies that complex interspecific interactions, implicitly represented by local limits to diversity, play a central macroecological role. Given its simplicity and apparent efficacy, it is plausible that with only minor changes to the model structure, the LSPOM could help dissect the drivers macroecological structures not considered here.

The LSPOM describes a process by which some species randomly gain territory at the expense of others. Because of the continuous arrival of new invaders, all species are eventually pushed to regional extinction and no species occupies very many patches. Thus, occupancy of more than one site is, in a sense, a fluctuation on the path leading from occupancy one for new invaders to occupancy zero, i.e. regional extinction. The near-extinct state, i.e. an occupancy of one, is therefore the most likely.

Our analysis does not necessarily imply that all sites surveyed are ecologically equivalent. Both OFD and metacommunity structure predicted by the LSPOM remain intact if the set of sites represented by the LSPOM is split into subsets representing different environmental conditions that are suitable for different kinds of species. The analogous argument was developed by Purves & Pacala (2005) and Chisholm & Pacala (2010) in the context of SADs. This observation makes it plausible why we previously found the log-series OFD predicted by the LSPOM in simulations of the Lotka-Volterra Metacommunity Model (LVMCM), where the abiotic environment varies from site to site. In the LVMCM the community at each site is represented by a Lotka-Volterra competition model and sites are linked through a planar dispersal network. This model has been shown to reproduce a variety of spatial and temporal macroecological patterns (O’Sullivan *et al.*, 2019, 2021). In contrast to the LSPOM, regulation of local species richness in the LVMCM is not an explicit constraint but an emergent phenomenon resulting from local ecological structural instability (Rossberg, 2013). The LVMCM has also shown how processes entirely intrinsic to the metacommunity can lead to local colonisation-extinction dynamics (O’Sullivan *et al.*, 2021) in a way that is closely analogous to the stochastic colonisations and extinctions that occur in patch-occupancy type models. Thus the LSPOM can be seen as an analytically tractable abstraction of the more complex LVMCM. For further discussion on the relationship between the two models see Supplementary information Sec. S8.

We caution that OFD determined according to substantially different protocols may not be readily explained by our theory. Storch & Šizling (2002), for example, studied an OFD that differed from ours by describing a group of species (birds) that is considerably more mobile, less species rich, and numerically less abundant. Their OFD quantified presence/absence in large, adjacent patches of 12 × 11 km extent, covering a wide range of habitat types of varying prevalence. Storch & Šizling (2002) could explain their OFD by a simple model combining habitat prevalence and species abundance information. With their study design, patch dynamics of the type considered here might indeed not arise or not be captured by the OFD. In particular, Storch & Šizling’s OFD is bimodal, while bimodality in the LSPOM steady state appears impossible, as it would require that some species never get replaced by others in any of the patches.

At finer spatial resolution, bimodality in the OFD has also been detected. Recent studies have reported qualitative temporal consistency in bimodal OFDs for assemblages of functionally similar species (Suhonen & Jokimäki, 2019; Suhonen, 2021). In our study systems bimodality is extremely rare, in particular after filtering out demonstrably limited samples. This is consistent with previous studies concluding that bimodality can plausibly arise artificially in studies on spatial scales small relative to typical species range sizes (van Rensburg *et al.*, 2000; Bossuyt *et al.*, 2004; Heatherly *et al.*, 2007).

Finally, in recent years evidence has been found of intrinsic biodiversity regulation, detectable as temporal invariance or stationarity in local community species richness (e.g. Brown *et al.* 2000; Parody *et al.* 2001; Magurran & Henderson 2010; Magurran *et al.* 2015). No systematic changes in local richness has been found at the global scale (Dornelas *et al.*, 2014; Gotelli *et al.*, 2017; Vellend *et al.*, 2017), though latitudinal and between-biome differences in dynamics have been shown (Blowes *et al.*, 2019). Our study builds on these important results, showing how such intrinsic constraints can propagate up to the metacommunity scale and ultimately determine macroecological structures such as the OFD.

## Footnotes

1 https://environment.data.gov.uk/ecology-fish/downloads/

2 I: https://environment.data.gov.uk/portalstg/sharing/rest/content/items/8aeb2f4f89304834b2d3195622bb47bb/data

3 P: https://environment.data.gov.uk/portalstg/sharing/rest/content/items/136453e009eb4f2489a8d1a15b610497/data

4 D: https://environment.data.gov.uk/portalstg/sharing/rest/content/items/1fc5164bcadf4f219ed34cc5d251c741/data

5 https://environment.data.gov.uk/catchment-planning/

6 https://data.gov.uk/dataset/1a494e3e-e414-456c-9c2e-ca367a2945b6/wfd-surface-water-management-catchments-cycle-2

## References

Álvarez-Noriega, M., Burgess, S.C., Byers, J.E., Pringle, J.M., Wares, J.P. & Marshall, D.J. (2020). Global biogeography of marine dispersal potential. Nature Ecology & Evolution, 4, 1196–1203.

Astorga, A., Death, R., Death, F., Paavola, R., Chakraborty, M. & Muotka, T. (2014). Habitat heterogeneity drives the geographical distribution of beta diversity: the case of New Zealand stream invertebrates. Ecology and Evolution, 4, 2693–2702.

Belmaker, J., Sekercioglu, C.H. & Jetz, W. (2012). Global patterns of specialization and coexistence in bird assemblages. Journal of Biogeography, 39, 193–203.

Blowes, S.A., Supp, S.R., Antão, L.H., Bates, A., Bruelheide, H., Chase, J.M., Moyes, F., Magurran, A., McGill, B., Myers-Smith, I.H. et al. (2019). The geography of biodiversity change in marine and terrestrial assemblages. Science, 366, 339–345.

Böhning-Gaese, K., Caprano, T., Ewijk, K.v. & Veith, M. (2006). Range size: disentangling current traits and phylogenetic and biogeographic factors. The American Naturalist, 167, 555–567.

Boieiro, M., Matthews, T., Rego, C., Crespo, L., Aguiar, C., Cardoso, P., Rigal, F., Silva, I., Pereira, F., Borges, P. & Serrano, A. (2018). A comparative analysis of terrestrial arthropod assemblages from a relict forest unveils historical extinctions and colonization differences between two oceanic islands. PloS One, 13, e0195492.

Bossuyt, B., Honnay, O. & Hermy, M. (2004). Scale-dependent frequency distributions of plant species in dune slacks: Dispersal and niche limitation. Journal of Vegetation Science, 15, 323–330.

Boswell, M.T. & Patil, G.P. (1971). Chance mechanisms generating the logarithmic series distribution used in the analysis of number of species and individuals. In: Statistical Ecology, Volume I, Spatial Patterns and Statistical Distributions (eds. Patil, G.P., Pielou, E.C. & Waters, W.E.). Pennsylvania State University Press, University Park, PA, pp. 99–130.

Boulangeat, I., Lavergne, S., Van Es, J., Garraud, L. & Thuiller, W. (2012). Niche breadth, rarity and ecological characteristics within a regional flora spanning large environmental gradients. Journal of Biogeography, 39, 204–214.

Bray, J.R. & Curtis, J.T. (1957). An ordination of the upland forest communities of southern Wisconsin. Ecological Monographs, 27, 326–349.

Brown, J.H. (1984). On the relationship between abundance and distribution of species. The American Naturalist, 124, 255–279.

Brown, J.H., Ernest, S.M., Parody, J.M. & Haskell, J.P. (2000). Regulation of diversity: maintenance of species richness in changing environments. Oecologia, 126, 321–332.

Brown, J.H., Mehlman, D.W. & Stevens, G.C. (1995). Spatial variation in abundance. Ecology, 76, 2028–2043.

Capurucho, J.a.M., Ashley, M.V., Tsuru, B.R., Cooper, J.C. & Bates, J.M. (2020). Dispersal ability correlates with range size in Amazonian habitat-restricted birds. Proceedings of the Royal Society B, 287, 20201450.

Carscadden, K.A., Emery, N.C., Arnillas, C.A., Cadotte, M.W., Afkhami, M.E., Gravel, D., Livingstone, S.W. & Wiens, J.J. (2020). Niche breadth: causes and consequences for ecology, evolution, and conservation. The Quarterly Review of Biology, 95, 179–214.

Chen, S.C., Tamme, R., Thomson, F.J. & Moles, A.T. (2019). Seeds tend to disperse further in the tropics. Ecology Letters, 22, 954–961.

Chisholm, R.A. & Pacala, S.W. (2010). Niche and neutral models predict asymptotically equivalent species abundance distributions in high-diversity ecological communities. Proceedings of the National Academy of Sciences, 107, 15821–15825.

Clark, C.W. & Rosenzweig, M.L. (1994). Extinction and colonization processes: parameter estimates from sporadic surveys. The American Naturalist, 143, 583–596.

Cleveland, W.S. & Devlin, S.J. (1988). Locally weighted regression: an approach to regression analysis by local fitting. Journal of the American Statistical Association, 83, 596–610.

Collins, S.L. & Glenn, S.M. (1997). Effects of organismal and distance scaling on analysis of species distribution and abundance. Ecological Applications, 7, 543–551.

Conover, W.J. (1999). Practical Non-parametric Statistics. vol. 350. john wiley & sons.

Day, J.R. & Possingham, H.P. (1995). A stochastic metapopulation model with variability in patch size and position. Theoretical Population Biology, 48, 333–360.

Delignette-Muller, M.L. & Dutang, C. (2015). fitdistrplus: An R package for fitting distributions. Journal of Statistical Software, 64, 1–34.

Denelle, P., Violle, C., Consortium, D. & Munoz, F. (2020). Generalist plants are more competitive and more functionally similar to each other than specialist plants: insights from network analyses. Journal of Biogeography, 47, 1922–1933.

Dornelas, M., Gotelli, N.J., McGill, B., Shimadzu, H., Moyes, F., Sievers, C. & Magurran, A.E. (2014). Assemblage time series reveal biodiversity change but not systematic loss. Science, 344, 296–299.

Drossel, B., Higgs, P.G. & McKane, A.J. (2001). The influence of predator–prey population dynamics on the long-term evolution of food web structure. Journal of Theoretical Biology, 208, 91–107.

Fuller, W.A. (2009). Introduction to Statistical Time Series. vol. 428. John Wiley & Sons.

Gaston, K.J. & He, F. (2002). The distribution of species range size: a stochastic process. Proceedings of the Royal Society of London. Series B: Biological Sciences, 269, 1079–1086.

Gotelli, N.J., Shimadzu, H., Dornelas, M., McGill, B., Moyes, F. & Magurran, A.E. (2017). Community-level regulation of temporal trends in biodiversity. Science Advances, 3, e1700315.

Hanski, I. (1982). Dynamics of regional distribution: the core and satellite species hypothesis. Oikos, pp. 210–221.

Harmon, L.J. & Harrison, S. (2015). Species diversity is dynamic and unbounded at local and continental scales. The American Naturalist, 185, 584–593.

Heatherly, T., Whiles, M., Gibson, D., Collins, S., Huryn, A., Jackson, J. & Palmer, M. (2007). Stream insect occupancy-frequency patterns and metapopulation structure. Oecologia, 151, 313–321.

Heino, J. (2015). Deconstructing occupancy frequency distributions in stream insects: effects of body size and niche characteristics in different geographical regions. Ecological Entomology, 40, 491–499.

Hubbell, S. (2001). The Unified Neutral Theory of Biodiversity and Biogeography (Prin. Princeton University Press, Princeton, NJ.

Jaccard, P. (1912). The distribution of the flora in the alpine zone. 1. New Phytologist, 11, 37–50.

Kendall, D.G. (1948a). On some modes of population growth leading to R A Fisher’s logarithmic series distribution. Biometrika, 35, 6–15.

Kendall, M.G. (1948b). Rank correlation methods.

Kennedy, M.P., Lang, P., Grimaldo, J.T., Martins, S.V., Bruce, A., Moore, I., Taubert, R., Macleod-Nolan, C., McWaters, S., Briggs, J. et al. (2017). Niche-breadth of freshwater macrophytes occurring in tropical southern African rivers predicts species global latitudinal range. Aquatic Botany, 136, 21–30.

Kwiatkowski, D., Phillips, P.C., Schmidt, P. & Shin, Y. (1992). Testing the null hypothesis of stationarity against the alternative of a unit root: How sure are we that economic time series have a unit root? Journal of Econometrics, 54, 159–178.

Laube, I., Korntheuer, H., Schwager, M., Trautmann, S., Rahbek, C. & Böhning-Gaese, K. (2013). Towards a more mechanistic understanding of traits and range sizes. Global Ecology and Biogeography, 22, 233–241.

Lester, S.E., Ruttenberg, B.I., Gaines, S.D. & Kinlan, B.P. (2007). The relationship between dispersal ability and geographic range size. Ecology Letters, 10, 745–758.

Levins, R. (1969). Some demographic and genetic consequences of environmental heterogeneity for biological control. American Entomologist, 15, 237–240.

Luo, B., Santana, S.E., Pang, Y., Wang, M., Xiao, Y. & Feng, J. (2019). Wing morphology predicts geographic range size in vespertilionid bats. Scientific Reports, 9, 1–6.

Magurran, A.E., Dornelas, M., Moyes, F., Gotelli, N.J. & McGill, B. (2015). Rapid biotic homogenization of marine fish assemblages. Nature Communications, 6, 1–5.

Magurran, A.E. & Henderson, P.A. (2010). Temporal turnover and the maintenance of diversity in ecological assemblages. Philosophical Transactions of the Royal Society B: Biological Sciences, 365, 3611–3620.

Mann, H.B. (1945). Non-parametric tests against trend. Econometrica: Journal of the Econometric Society, pp. 245–259.

McCaslin, H.M., Caughlin, T.T. & Heath, J.A. (2020). Long-distance natal dispersal is relatively frequent and correlated with environmental factors in a widespread raptor. Journal of Animal Ecology, 89, 2077–2088.

McGeoch, M.A. & Gaston, K.J. (2002). Occupancy frequency distributions: patterns, artefacts and mechanisms. Biological Reviews, 77, 311–331.

McGill, B. (2003). Strong and weak tests of macroecological theory. Oikos, 102, 679–685.

McGill, B.J., Etienne, R.S., Gray, J.S., Alonso, D., Anderson, M.J., Benecha, H.K., Dornelas, M., Enquist, B.J., Green, J.L., He, F. et al. (2007). Species abundance distributions: moving beyond single prediction theories to integration within an ecological framework. Ecology Letters, 10, 995–1015.

Moritz, S. & Bartz-Beielstein, T. (2017). imputeTS: Time Series Missing Value Imputation in R. The R Journal, 9, 207–218.

Mykrä, H. & Heino, J. (2017). Decreased habitat specialization in macroinvertebrate assemblages in anthropogenically disturbed streams. Ecological Complexity, 31, 181–188.

O’Sullivan, J.D., Knell, R.J. & Rossberg, A.G. (2019). Metacommunity-scale biodiversity regulation and the self-organised emergence of macroecological patterns. Ecology letters, 22, 1428–1438.

O’Sullivan, J.D., Terry, J.C.D. & Rossberg, A.G. (2021). Intrinsic ecological dynamics drive biodiversity turnover in model metacommunities. Nature Communications, 12, 1–11.

O’Sullivan, J.D., Terry, J.C.D. & Rossberg, A.G. (2022). Community composition exceeds area as a predictor of long-term conservation value. In preparation.

Pagel, M.D., May, R.M. & Collie, A.R. (1991). Ecological aspects of the geographical distribution and diversity of mammalian species. The American Naturalist, 137, 791–815.

Parody, J.M., Cuthbert, F.J. & Decker, E.H. (2001). The effect of 50 years of landscape change on species richness and community composition. Global Ecology and Biogeography, 10, 305–313.

Pawar, S. (2009). Community assembly, stability and signatures of dynamical constraints on food web structure. Journal of Theoretical Biology, 259, 601–612.

Pearson, K. (1900). On the criterion that a given system of deviations from the probable in the case of a correlated system of variables is such that it can be reasonably supposed to have arisen from random sampling. The London, Edinburgh, and Dublin Philosophical Magazine and Journal of Science, 50, 157–175.

Pianka, E.R. (1966). Latitudinal gradients in species diversity: a review of concepts. The American Naturalist, 100, 33–46.

Pielou, E.C. (1977). Mathematical ecology. Wiley.

Pigolotti, S., Flammini, A., Marsili, M. & Maritan, A. (2005). Species lifetime distribution for simple models of ecologies. Proceedings of the National Academy of Sciences, 102, 15747–15751.

Preston, F.W. (1948). The commonness, and rarity, of species. Ecology, 29, 254–283.

Pueyo, S., He, F. & Zillio, T. (2007). The maximum entropy formalism and the idiosyncratic theory of biodiversity. Ecology Letters, 10, 1017–1028.

Purves, D.W. & Pacala, S.W. (2005). Ecological drift in niche-structured communities: neutral pattern does not imply neutral process. In: Biotic interactions in the tropics: Their Role in the Maintenance of Species Diversity (eds. Burslem, D., Pinard, M. & Hartley, S.). Cambridge University Press, pp. 107–138.

Rabosky, D.L. & Hurlbert, A.H. (2015). Species richness at continental scales is dominated by ecological limits. The American Naturalist, 185, 572–583.

Raunkiaer, C. et al. (1934). The life forms of plants and statistical plant geography; being the collected papers of C. Raunkiaer. The life forms of plants and statistical plant geography; being the collected papers of C. Raunkiaer

Renner, S., Dalzochio, M.S., Périco, E., Sahlén, G. & Suhonen, J. (2020). Odonate species occupancy frequency distribution and abundance–occupancy relationship patterns in temporal and permanent water bodies in a subtropical area. Ecology and Evolution, 10, 7525–7536.

van Rensburg, B.J., McGeoch, M.A., Matthews, W., Chown, S.L. & van Jaarsveld, A.S. (2000). Testing generalities in the shape of patch occupancy frequency distributions. Ecology, 81, 3163–3177.

Rossberg, A.G. (2013). Food webs and biodiversity: foundations, models, data. John Wiley & Sons.

Ruggiero, A. (1994). Latitudinal correlates of the sizes of mammalian geographical ranges in South America. Journal of Biogeography, pp. 545–559.

Sarremejane, R., Cid, N., Stubbington, R., Datry, T., Alp, M., Cañedo Argüelles, M., Cordero-Rivera, A., Csabai, Z., Gutiérrez-Cánovas, C., Heino, J. et al. (2020). DISPERSE, a trait database to assess the dispersal potential of European aquatic macroinvertebrates. Scientific Data, 7, 1–9.

Schloss, C.A., Nuñez, T.A. & Lawler, J.J. (2012). Dispersal will limit ability of mammals to track climate change in the Western Hemisphere. Proceedings of the National Academy of Sciences, 109, 8606–8611.

Sen, P.K. (1968). Estimates of the regression coefficient based on Kendall’s tau. Journal of the American Statistical Association, 63, 1379–1389.

Sheard, C., Neate-Clegg, M.H., Alioravainen, N., Jones, S.E., Vincent, C., MacGregor, H.E., Bregman, T.P., Claramunt, S. & Tobias, J.A. (2020). Ecological drivers of global gradients in avian dispersal inferred from wing morphology. Nature Communications, 11, 1–9.

Slatyer, R.A., Hirst, M. & Sexton, J.P. (2013). Niche breadth predicts geographical range size: a general ecological pattern. Ecology letters, 16, 1104–1114.

Stevens, G.C. (1989). The latitudinal gradient in geographical range: how so many species coexist in the tropics. The American Naturalist, 133, 240–256.

Storch, D. & Šizling, A.L. (2002). Patterns of commonness and rarity in central European birds: reliability of the core-satellite hypothesis within a large scale. Ecography, 25, 405–416.

Suhonen, J. (2021). Spatial and temporal changes in occupancy frequency distribution patterns of freshwater macrophytes in Finland. Ecology and Evolution.

Suhonen, J. & Jokimäki, J. (2019). Temporally stable species occupancy frequency distribution and abundance–occupancy relationship patterns in urban wintering bird assemblages. Frontiers in Ecology and Evolution, 7, 129.

Takashina, N., Plank, M.J., Jenkins, C.N. & Economo, E.P. (2022). Species-range-size distributions: Integrating the effects of speciation, transformation, and extinction. Ecology and Evolution, 12, e8341.

Tamme, R., Götzenberger, L., Zobel, M., Bullock, J.M., Hooftman, D.A., Kaasik, A. & Pärtel, M. (2014). Predicting species’ maximum dispersal distances from simple plant traits. Ecology, 95, 505–513.

Theil, H. (1950). A rank-invariant method of linear and polynomial regression analysis. Indagationes Mathematicae, 12, 173.

Tokeshi, M. (1992). Dynamics of distribution in animal communities: theory and analysis. Researches on Population Ecology, 34, 249–273.

Tukey, J.W. (1977). Box-and-whisker plots. Exploratory Data Analysis, pp. 39–43.

Vellend, M., Dornelas, M., Baeten, L., Beauséjour, R., Brown, C.D., De Frenne, P., Elmendorf, S.C., Gotelli, N.J., Moyes, F., Myers-Smith, I.H. et al. (2017). Estimates of local biodiversity change over time stand up to scrutiny. Ecology, 98, 583–590.

Volkov, I., Banavar, J.R., Hubbell, S.P. & Maritan, A. (2003). Neutral theory and relative species abundance in ecology. Nature, 424, 1035–1037.

Watterson, G.A. (1974). Models for the logarithmic species abundance distributions. Theoretical Population Biology, 6, 217–250.

Wayman, J.P., Sadler, J.P., Pugh, T.A., Martin, T.E., Tobias, J.A. & Matthews, T.J. (2022). Assessing taxonomic and functional change in British breeding bird assemblages over time. Global Ecology and Biogeography, 31, 925–939.

Williams, C. (1947). The logarithmic series and its application to biological problems. Journal of Ecology, 34, 253–272.

Yoshida, K. (2003). Evolutionary dynamics of species diversity in an interaction web system. Ecological Modelling, 163, 131–143.

