## Supplementary Information for "Temporally robust occupancy frequency distributions in riverine metacommunities explained by local biodiversity regulation"

#### S1 Filtering the EA databases

The survey data included $4138$, $2952$, and $2816$ unique water body identifiers in the macroinvertebrate (I), macrophyte (P) and diatom (D) datasets, respectively. Of these, however, $597$, $297$, and $339$ waterbodies, respectively, had no available management catchment identifiers and these were dropped. The three datasets represent distinct observation programs and the sites surveyed in a given catchment year don't necessarily correspond between programs. Hence the following filtering steps were applied to each of the datasets independently. Since a clear trend in numbers of sites surveyed and taxa observed was found in the earliest years of the surveys, we excluded all observations prior to $1990$ from the analysis, leaving catchment OFD time series spanning around $30$ years. We constrained our analysis to catchment-years in which a minimum of $10$ taxa were detected, and a minimum of $5$ sites were sampled. This helped to reduce small-sample-size effects in the estimated moments.

Since we are particularly interested in the macroscopic shape of the OFD best represented by its skewness, we used a rarefaction method to filter the dataset of catchment-years for which the estimated skewness of the OFD was unlikely to be biologically representative. For each catchment-year, we constructed a binary taxon-by-site table representing the presence-absence of taxa at each site in that year's survey. We then generated $1000$ independent randomisations of the taxon-by-site table by randomly subsampling a random number of taxa between $1$ and the total observed richness for the catchment-year. For these random subsets of the empirical community we computed the skewness of the OFD. Then we numerically estimated the first derivative of the skewness estimate as a function of the size of the random subsample. Catchment-years for which the average absolute value of the first derivative evaluated between $90\%$ and $100\%$ of the observed richness was greater than $10^{-2}$ were considered to have failed to converge and dropped from the analysis (Fig. S1), ensuring representative skewness estimates despite unavoidable type II errors in the surveys. Finally, once these catchment-years were excluded, we dropped any catchment from the analysis for which the remaining time series included less than $8$ years of data. For catchments retained following filtering but for which single catchment-years were missing, we used a linear interpolation method prior to time series analyses to handle missing data points in the time series (Moritz & Bartz-Beielstein, 2017).

Prior to filtering, the database included $2708$, $1625$, $1120$ catchment-years (unique OFDs) split between $97$, $92$, and $90$ catchments for the I, P and D datasets respectively. Following these various filtering steps, we ended up with $1804$, $731$ and $588$ catchment-years in $83$, $52$ and $48$ catchments, respectively.

#### S2 Simulating the LSPOM

For illustrative purposes it can be informative to numerically simulate a LSPOM metacommunity in which stochastic colonisation and extinction processes occur as described in Box 1. R code for the simple simulation is available in the accompanying Github repository, here we outline the mechanics of the simulation.

Each of the $N$ sites in the spatially implicit metacommunity hosts $\alpha$ local populations. The state of the metacommunity is represented by an $N$ by $\alpha$ occupancy table with each element indicating the identity of the species to which a given local population belongs. In each iteration of the stochastic patch occupancy model, the occupancy table is updated in response to the arrival of a single new invader into the metacommunity from outside as well as stochastic colonisation-extinction dynamics internal to the assemblage.

Each newly introduced species invades a single site—the number of sites initially occupied, and hence the invasion fitness of each species, is identical. The site into which the species invades is randomly sampled with a probability of $\alpha_i/Z$ where $i$ is a site index and where $Z = \sum_i^N \alpha_i$ is the total number of local populations. For simplicity $\alpha_i$ are identical for all sites. Since local species richness is strictly limited, each colonisation produces a single local extirpation event. The probability of extirpation following local colonisation is identical for all species—the identity of the local resident excluded is randomly sampled with a probability of $1/\alpha_i$. Following each regional invasion, within-metacommunity colonisation extinction occurs. Each local population expands its range into a single new site with a probability $m/Z$. The colonised site and the identity of the extirpated resident are randomly sampled with the same probability as in the case of regional-scale invasions. In the numerical simulation, the internal colonisation-extinctions occur sequentially.

Fig. S4 shows the steady state OFD that emerges in such LSPOM simulations. The points represent the average richness in each occupancy class across $10^3$ replicate simulations. In each case, $N = 50$ sites, each hosting $\alpha_i = 10$ populations, and $10^4$ iterations were modelled. The shape of the emergent OFD is controlled by the mixing rate parameter $m$ which in Fig. S4 we set to 2.5, 5 and 10.

#### S3 Elementary derivation of the log-series steady state OFD

The mathematics underlying our model is well understood from the analogous model for the SAD (e.g. Volkov *et al.* 2003). Here we provide a brief elementary derivation formulated in the OFD context.

First, let $X_n(t)$ denote the time-dependent expectation value of the number of species occupying $n$ patches at a given time $t$. Recall that $N$ is the number of patches, $\alpha_i$ the number of populations in each patch, and $Z = \sum_i^N \alpha_i$

the total number of populations. In order to find the long-term equilibrium richness in each occupancy class we must consider the probability that these expectation values change through time. If each $n$ is understood as a 'compartment' containing $X_n(t)$ species, transitions into or out of the $n$th compartment can occur in the following five ways:

(i) $X_n(t)$ increases by one due to the transition of a species out of the $(n-1)$th occupancy class—stochastic range expansion.

(ii) $X_n(t)$ increases by one due to the transition of a species out of the $(n+1)$th occupancy class—range reduction due to local extirpation.

(iii) $X_n(t)$ decreases by one due to the transition of a species from the $n$th to the $(n+1)$th occupancy class—stochastic range expansion.

(iv) $X_n(t)$ decreases by one due to the transition of a species from the $n$th to the $(n-1)$th occupancy class—range reduction due to local extirpation.

(v) For the $n = 1$ case only, $X_1(t)$ increases by one due to invasion into the metacommunity from outside.

Note that in this scheme we assume that $N$ is much greater than typical species occupancies such that we can disregard the case that a species colonises a patch in which it is already present. In the numerical realisation of the model, when choosing patches to colonise at random, invocation of this constraint would be rare if, for a given species, there are many more unoccupied than occupied patches.

Now let's consider the rates (probabilities) of each of these five transitions. As described in Box 1, by construction any extant population colonises an additional patch at a rate of $m/Z$ per unit time. Thus, (i) and (iii) occur at a mean rate of $m/Z$. To determine the rate of extirpation of species from patches which are saturated with respect to the number of populations they can sustain, i.e. of processes (ii) and (iv), consider that a the population of a given species resident at a given patch $i$ can get extirpated as a result of colonisation of that patch by any of the $Z - \alpha_i \approx Z$ populations from outside that patch. On the other hand, a given population from a given patch $j$ extirpates any population in any other patch with the same probability, so it extirpates that population in patch $i$ with probability $1/(Z-\alpha_j) \approx 1/Z$. Recalling that each population colonises other patches at rate $m/Z$, the rate of extirpation of the focal population in patch $i$ due to colonisation from any other patches is therefore approximately $Z \times (1/Z) \times (m/Z) = m/Z$.

We can now write down a differential equation for the rate of change of $X_n = X_n(t)$ due to the five processes explained above. For $n > 1$

$$\frac{dX_n}{dt} = \overbrace{\frac{m}{Z}(n-1)X_{n-1}}^{\text{(i)}} + \overbrace{\frac{m+1}{Z}(n+1)X_{n+1}}^{\text{(ii)}} - \overbrace{\frac{m}{Z}nX_n}^{\text{(iii)}} - \overbrace{\frac{m+1}{Z}nX_n}^{\text{(iv)}} \tag{S1}$$

Regional invaders can only transition into the 1st occupancy class. Thus for $X_1$

$$\frac{dX_1}{dt} = \overbrace{1}^{\text{(v)}} + \overbrace{\frac{m+1}{Z}2X_2}^{\text{(ii)}} - \overbrace{\frac{m}{Z}X_1}^{\text{(iii)}} - \overbrace{\frac{m+1}{Z}X_1}^{\text{(iv)}} \tag{S2}$$

In the steady state, $dX_n/dt = 0$ for both differential equations. Substituting for $X_n$ the log-normal OFD in Eq. 1 into Eqs. (S1) and (S2), one can see that it solves the equation; in both cases, the first term balances the last and the second term the third. While the equilibrium condition permits other solutions, only Eq. 1 also satisfies the consistency condition that the expected total number of populations in the system equals $Z$, that is, assuming $N \gg m$,

$$\sum_{n=1}^{\infty} nX_n = Z. \tag{S3}$$

### S4 Estimation of false-negative and extirpation rates

To estimate both the mean false-negative and the mean extirpation rates of species in catchments, we determined for each pair $(y_1, y_2)$ of recorded years with $y_1 < y_2$ the proportion $\hat{p}(y_1, y_2)$, across all sites, of species observed at a site in $y_1$ that were also observed at that site in $y_2$.

A theoretical model for the expectation value of $\hat{p}(y_1, y_2)$ is given by

$$\begin{aligned} p(y_1, y_2) &= \omega \left\{ (1-d) \exp\left[ -\frac{e\,(y_2 - y_1)}{1-d} \right] + d \right\} \\ &= (\omega - \hat{d}) \exp[-\hat{e}\,(y_2 - y_1)] + \hat{d}, \end{aligned} \tag{S4}$$

where the parameter $\omega$ represents the probability of observing a species that is present at a site ($1 - \omega$ is the false-negative rate), $e$ is the site-level extirpation rate, $d$ is the unconditional probability that a randomly chosen species occupies a randomly chosen site, and we defined $\hat{d} = \omega d$ and $\hat{e} = e/(1-d)$. Equation (S4) describes the decline of the probability of re-observing a species at a site where it was previously observed. It derives from a continuous-time Markov model with transitions between an occupied and a vacant state at rates $e$ (extirpation) and $c$ (colonisation), which has a steady-state occupation probability of $d = c/(e+c)$ and a relaxation rate to the steady state of $\hat{e} = e + c$. For simplicity, the model underlying Equation (S4) disregards catchment-level species turnover, which would lead to a gradual decline of colonisation rate $c$ on average.

We estimated $\hat{d}$ directly from empirical site-occupancy tables as the mean number of species observed in sampled sites, averaged over all years, divided by the total number of species observed in a catchment, averaged over all years. Then we obtained $\omega$ and $\hat{e}$ from a weighted least-square fit of $\hat{p}(y_1, y_2)$ to Eq. (S4) over all pairs $(y_1, y_2)$. From the model fits we then obtained the extirpation rate as $e = \hat{e}(1 - \hat{d}/\omega)$. Occasionally, this fitting algorithm yielded estimates of $\omega$ slightly larger than one. We retained these values, despite them being theoretically impossible, to avoid biasing the overall estimation procedure.

The rationale for the least-square fit is as follows. Conceptually, we first construct a smooth curve for the empirical probability $\hat{p}(y_2 - y_1)$ of re-observation after a given time lag $y_2 - y_1$ by smoothing the empirical proportions $\hat{p}(y_1, y_2)$ of re-observed species over lags. The smoothing involves some kind of averaging, i.e. least-square fits (recall that the mean $\overline{x}$ of a set of values value $x_1, \ldots, x_n$ equals the value $x_\mathrm{c}$ that minimises the sum of square $\sum_i^n (x_i - x_\mathrm{c})^2$). Then we fit the right-hand side of Equation (S4) to $\hat{p}(y_2 - y_1)$. The intermediate step of constructing $\hat{p}(y_2 - y_1)$, however, is redundant. Instead, we directly perform a least-square fit of $\hat{p}(y_1, y_2)$ to $p(y_1, y_2)$, combining smoothing and fitting into a single step. As well as offering greater algorithmic simplicity, this approach has the advantage of naturally taking account of unevenness in the distribution of lags $y_2 - y_1$ (for example, there is more data for small than for large lags). When fitting the models, the weight for a pair $(y_1, y_2)$ was chosen as the geometric mean over years $y_1$ and $y_2$ of the sum of empirical species richness over all site (= the sum of empirical occupancy over all species). This weighting was chosen for the high robustness of fitted $\omega$ and $\hat{e}$ values it provided under removal of the data for any single year.

We also considered even more sophisticated methods that estimate observation, extirpation, and re-colonisation probability by fitting Hidden Markov Models of site occupancy directly to recorded sequences of observations and non-observations Clark & Rosenzweig (1994). However, we decided not to use such models out of concern that they would struggle with the high variability in sampling effort in our data set.

We found median extirpation rates of $0.05$, $0.04$ and, $0.11$ per year for I, P, and D metacommunities, respectively. Estimated average false-negative rates ranged from $(1 - \omega)$ in the range $0.17$-$0.51$, $0.21$-$0.47$, $0.0$-$0.54$

respectively. The distributions of these key rates are shown in Fig. S16.

#### S5 The effect of false negatives on log-series OFD

We compute the effect of false negatives on log-series OFD. The result is known since Williams (1947) (who attributes it to 'M. H. Quenouille') and Kendall (1948a), but neither provide a derivation. We provide it here for completeness and to illustrate its intuitive simplicity when formulated in terms of the mixing-rate parameter.

The derivation makes use of probability generating functions (PGFs). For any random variable attaining non-negative random values $n$, its PGF $G(z)$ is defined as the expectation value of $z^n$, i.e., $G(z) = \sum_n p_n z^n$, where $p_n$ is the probability of value $n$ and $z$ is a formal book-keeping variable.

Let

$$p_n = \frac{1}{n \ln(m+1)} \left( \frac{m}{m+1} \right)^n \tag{S5}$$

be the probability for a randomly sampled extant species to have occupancy $n$. The corresponding PGF is

$$G_{\text{OFD}}(z) = \sum_{n=1}^{\infty} p_n z^n = -\frac{\ln\left(1 - \frac{m\,z}{m+1}\right)}{\ln(m+1)} = \frac{\ln\left(\frac{m+1}{m+1-m\,z}\right)}{\ln(m+1)}. \tag{S6}$$

Furthermore, the PGF for a Bernoulli variable that is $1$ with probability $\omega$ and $0$ otherwise is:

$$G_{\text{BER}}(z) = (1-\omega) + \omega\, z. \tag{S7}$$

It follows directly from the definition of PGFs that the PGF for the sum of independent instances of a random variable is the product of their PGFs. Hence, for a species with occupancy $n$, the PGF for the number of sites at which it is observed is $[G_{\text{BER}}(z)]^n$ provided $(1-\omega)$ is the false-negative rate. If $n$ is a random variable with the log-series distribution given by Eq. (S5), the corresponding PGF is

$$G_{\text{OBS}}(z) = \sum_{n=1}^{\infty} p_n [G_{\text{BER}}(z)]^n = G_{\text{OFD}}[G_{\text{BER}}(z)]. \tag{S8}$$

From the definition of the PGF, it follows that the probability of not observing a randomly selected extant species at all is

$$P_{\text{miss}} = \lim_{z \to 0^+} G_{\text{OBS}}(z) = G_{\text{OFD}}(1-\omega) = \frac{\ln\left(\frac{m+1}{m\,\omega+1}\right)}{\ln(m+1)}. \tag{S9}$$

Since empirical OFD consider only those species that have been observed as extant, we need to convert $G_{\text{OBS}}(z)$ to the corresponding PGF conditional to observing a species at least once. This is obtained as

$$G_{\text{OFD,emp}}(z) = \frac{G_{\text{OBS}}(z) - P_{\text{miss}}}{1 - P_{\text{miss}}} = \frac{\ln\left(\frac{m\,\omega+1}{m\,\omega+1-m\omega\,z}\right)}{\ln(m\,\omega+1)}. \tag{S10}$$

Comparison with Eq. (S6) shows that non-zero false-negative rates modify log-series OFD by effectively replacing the parameter $m$ by $m\,\omega$.

From analogous arguments we conclude that an additional modification of the parameter $m$ would be required when the sites sampled are a subset of the 'actual number of sites' of the metacommunity. For completeness, if $q$ is the proportion of the metacommunity sampled, the estimated parameter $m$ should be replaced by the composite value $m\,\omega\,q$. Since $q$ cannot be estimated from the available data, we assume here that $q \approx 1$ and disregard this correction.

### S6 The probability of finding a species also in another subsample of patches of a metacommunity

Consider a large metacommunity of which a random subset of patches that corresponds to a proportion $q$ of the entire metacommunity has been sampled (precisely, local richness $\alpha_i$ summed over these patches is a proportion $q$ of the overall sum $Z$, see Box 1). Here we ask what the proportion $P_{\text{other}}$ of the species sampled in this subset will on average that is also be found in a second random subset of equal size. We focus on the particular case that the second subset is selected conditional to not overlapping with the first subset. If some overlap is permitted, the proportion from the first subet found also in the second subset of patches can only be larger on average. Hence, the value $P_{\text{other}}$ we compute represents a lower bound for the situation of arbitrary degrees of overlap.

We assume that the metacommunity has a log-series OFD and that species are otherwise randomly distirbuted over patches. The problem is then equivalent to the following: for a single species with occupancy probability distribution given by Eq. (S5), assign each of its populations at random to three different types, which correspond to the two distrinct subsamples of patches and the remaining set of patches: to Type 1 with probabiltiy $q$, to Type 2 with probability $q$, and otherwise, with probability $1 - 2q$, to Type 3. Denote by $p_{n,m,r}$ the probability that this results in there being $n$ population of Type 1, $m$ populations of Type 2, and $l$ populations of Type 3. Then ask what the probability $P_{\text{other}}$ is that there is at least one population of Type 2, conditional to there being at least one of Type 1.

Define a PGF corresponding to $p_{m,n,l}$ as

$$G_{\text{ODF3}}(x, y, w) = \sum_{m=0}^{\infty} \sum_{n=0}^{\infty} \sum_{l=0}^{\infty} p_{m,n,l} x^n \, y^m \, w^l, \tag{S11}$$

that is, the expectation value of $x^n y^m w^l$. By arguments analogous to those invoked in Supplementary Information S5, we obtain $G_{\text{ODF3}}(x, y, w)$ from $G_{\text{ODF}}(z)$, given by Eq. (S6), as

$$G_{\text{ODF3}}(x, y, w) = G_{\text{ODF}}(qx + qy + (1 - 2q)w)\,. \tag{S12}$$

The probability that none of the populations is of Type 1 is given by

$$P_{\text{miss1}} = G_{\text{ODF3}}(0, 1, 1) = G_{\text{ODF}}(1 - q), \tag{S13}$$

as is easily verified by evaluating Eq. (S11) with $x = 0$ and $y = w = 1$.

To condition the joint distribution of $m$, $n$, $l$ on $m \geq 1$, we need to remove the cases where $m = 0$ and then divide by $1 - P_{\text{miss1}}$ to re-establish normalisation. This leads to a corresponding PGF

$$G_{\text{ODF3+}}(x, y, w) = \frac{G_{\text{ODF3}}(x, y, w) - G_{\text{ODF3}}(0, y, w)}{1 - P_{\text{miss1}}}. \tag{S14}$$

From this we obtain the conditional probability that no patch is of Type 2 as $G_{\text{ODF3+}}(1, 0, 1)$ and conversely the conditional probability that one or more patches are of Type 2 as

$$
\begin{aligned}
P_{\text{other}} = 1 - G_{\text{ODF3}+}(1,0,1) &= 1 - \frac{G_{\text{ODF3}}(1,0,1) - G_{\text{ODF3}}(0,0,1)}{1 - P_{\text{miss1}}} \\
&= 1 - \frac{G_{\text{ODF}}(1-q) - G_{\text{ODF}}(1-2q)}{1 - G_{\text{ODF}}(1-q)} \\
&= \frac{1 - 2G_{\text{ODF}}(1-q) + G_{\text{ODF}}(1-2q)}{1 - G_{\text{ODF}}(1-q)} \\
&= \frac{2 - 2G_{\text{ODF}}(1-q) - 1 + G_{\text{ODF}}(1-2q)}{1 - G_{\text{ODF}}(1-q)} \\
&= 2 - \frac{1 - G_{\text{ODF}}(1-2q)}{1 - G_{\text{ODF}}(1-q)}.
\end{aligned} \tag{S15}
$$

Using Eq. (S6), this evaluates to

$$
P_{\text{other}} = 2 - \frac{\ln(1+2mq)}{\ln(1+mq)}. \tag{S16}
$$

There are situations, such as ours, where neither $m$ nor $q$ are known, but where the OFD in the first sampled subset of patches follows a log-series distribution with mixing rate parameter $\omega m_{\text{OFD}}$, with an observation probability $\omega$ that can be independently estimated. In Supplementary Information S5, we argued that this fitted mixing rate corresponds to $m\omega q$. Dividing by $\omega$ to correct for false negatives, we therefore obtain $m_{\text{OFD}} = mq$, the quantity entering Eq. (S16). This allows us to estimate

$$
P_{\text{other}} = 2 - \frac{\ln(1+2m_{\text{OFD}})}{\ln(1+m_{\text{OFD}})} \tag{S17}
$$

without any knowlege of $q$.

### S7 Comparing LSPOM colonisation rate to observed turnover

Many different theories have been put forward for how and why log-series distributions arise in ecology, mostly in the context of SAD (Kendall 1948a; Watterson 1974; Boswell & Patil 1971; Pueyo *et al.* 2007 but see also Gaston & He 2002). Many of these theories are directly transferable to OFD. When including other families of right-skewed distributions, which can be empirically indistinguishable, the variety of explanations is even larger (McGill *et al.*, 2007; Takashina *et al.*, 2022). The need for "moving beyond single prediction theories" to identify the correct explanations of these distributions therefore arises equally for OFD as it does for SAD (McGill *et al.*,

2007).

Here we develop a method to compute alternative estimates of the mixing rate $m$ based not on the spatial structure of the metacommunity represented by the OFD, but instead on the temporal turnover rate. This method builds upon previous work on the Unified Neutral Theory of Biodiversity (UNTB) (Hubbell, 2001), extending it into the patch occupancy framework in which numbers of occupied patches are modelled instead of numbers of individuals. The analogues of speciations, births, and deaths in the UNTB are, respectively, regional invasions into the metacommunity, colonisations of new sites by members of the metacommunity, and local extirpations resulting from colonisations. This formal analogy allows us to adapt analytical insights developed for neutral individual-based community models to the patch occupancy framework. The aim of this method is to provide greater support to the LSPOM representation by showing quantitative consistency in the mixing rate estimates. If this could be shown we would be in a position to confirm that the LSPOM quantitatively captures both the shape of the OFD *and* the composite demographic rates summarised in the mixing rate parameter, $m$. Variation in site selection between years means the method cannot be applied with a high degree of confidence. We report the method here as a demonstration of the concept and aim to apply this or a similar analysis to alternative datasets in future to further probe the range of biological processes plausibly described by the neutral patch-dynamics framework.

From the UNTB and its various extensions, the probability distribution of the time to regional extinction of a randomly chosen species (called regional persistence time) is a known function of only two parameters (Pigolotti *et al.*, 2005): the mixing rate $m$ and the rate of local extirpation rate $e$. After estimating $e$ directly from observed local species turnover (see Sec. S4), it is possible to obtain a second, independent estimate of $m$ from fitting this function to the observed regional persistence time distribution. To achieve this, one must determine for each pair $(y_1, y_2)$ of recorded years with $y_1 < y_2$ the proportion $P(y_1, y_2)$ of those species regionally extant in $y_1$ that were also extant in $y_2$. One must then compute a weighted least-square fit of the model

$$P(y_1, y_2) = \Omega \left\{ 1 - \frac{1}{\log(1-\alpha)} \log\left[ \frac{(1-\alpha)\exp\left[(1-\alpha)\, e\, t\right]}{\exp\left[(1-\alpha)\, e\, t\right] - \alpha} \right] \right\} \tag{S18}$$

over all these pairs $(y_1, y_2)$, setting $t = y_2 - y_1$. The fitting parameter $\Omega$ represents the probability of regionally observing a species that is regionally extant,

$$\alpha = \frac{m}{m+1}, \tag{S19}$$

and the expression in braces is the theoretical complementary cumulative regional persistence time distribution (see Eq. 10 of Pigolotti *et al.*, 2005). We used the same weights as for the determination of $\omega$ and $e$. Thus, by analysis of the catchment-scale persistence time one can compute an alternative estimate of $m$, denoted $m_{\mathrm{TS}}$.

In Fig. S17 we report the mixing rates $m_{\mathrm{OFD}}$ and $m_{\mathrm{TS}}$ estimated based on the LSPOM log-series and time series analyses respectively. The ratio between the median estimates computed from OFD and turnover analyses Med $(\hat{m}_{\mathrm{OFD}})$/Med $(\hat{m}_{\mathrm{TS}})$ was in the range 4.12-8.22 depending on taxon. Thus, with the current data, we cannot conclusively show that the demographic rates implicit in the log-series model of empirical OFDs quantitatively coincide with plausible alternative estimates, we find that the differences are nonetheless strikingly small. It is plausible that a more systematic sampling procedure and better understanding of the sampling coverage $q$ (currently overlooked in our estimates) would reveal greater quantiative agreement. The method laid out here, or indeed some analogous union of the LSPOM and the UNTB, may offer an avenue for further validation of the biological plausibility of the log-series OFD in future work.

### S8 A meeting of models: a brief comparison of the LVMCM and the LSPOM

The LVMCM is a spatially explicit, non-neutral, population-dynamical metacommunity model. It describes the time evolution of all species' population biomasses as a function of local environmental conditions, ecological interactions and dispersal. In contrast, the LSPOM is a spatially and mechanistically implicit, ecologically neutral, probabilistic patch occupancy model. Despite these seemingly irreconcilable structural differences, the LSPOM captures both the dominant processes and the emergent steady-state biodiversity distribution of the LVMCM (in a large part of its parameter space). Crucially, its parametric simplicity means the LSPOM can be fitted directly to survey data. Elsewhere (O'Sullivan *et al.*, 2022) we developed a method for extracting spatial and environmental distributions and local species richness records from empirical observations and inputting these directly into LVMCM simulations. Parameters could then be tuned to match the estimated LSPOM mixing rates of the observation, thus reproducing the empirical macroecology. Thus, we also propose the LSPOM as a potential bridge between empirical dataset and more complex models such as the LVMCM which can not be easily parameterised empirically.

### S9 Supplementary figures

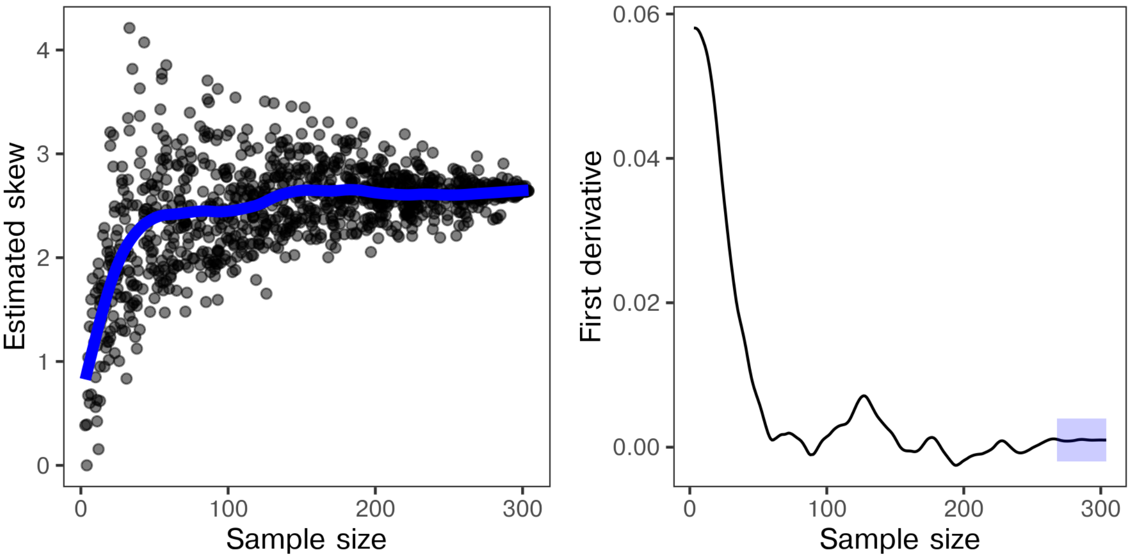


Figure S1: **Skewness of the OFD computed after randomly subsampling species to various 'depths' (number of species).** The average of the first derivative evaluated over the last $10\%$ of the sub-sample depth (blue box) is taken to represent the degree of convergence in the estimate. Catchment-years for which the evaluated first derivative, averaged over this region of the randomisation experiment, had an absolute value of greater than $10^{-2}$ were considered to have failed to converge. In such cases we concluded the sampling effort for the given year was insufficient to accurately describe the shape of the OFD and the data were dropped from the down-stream analyses.

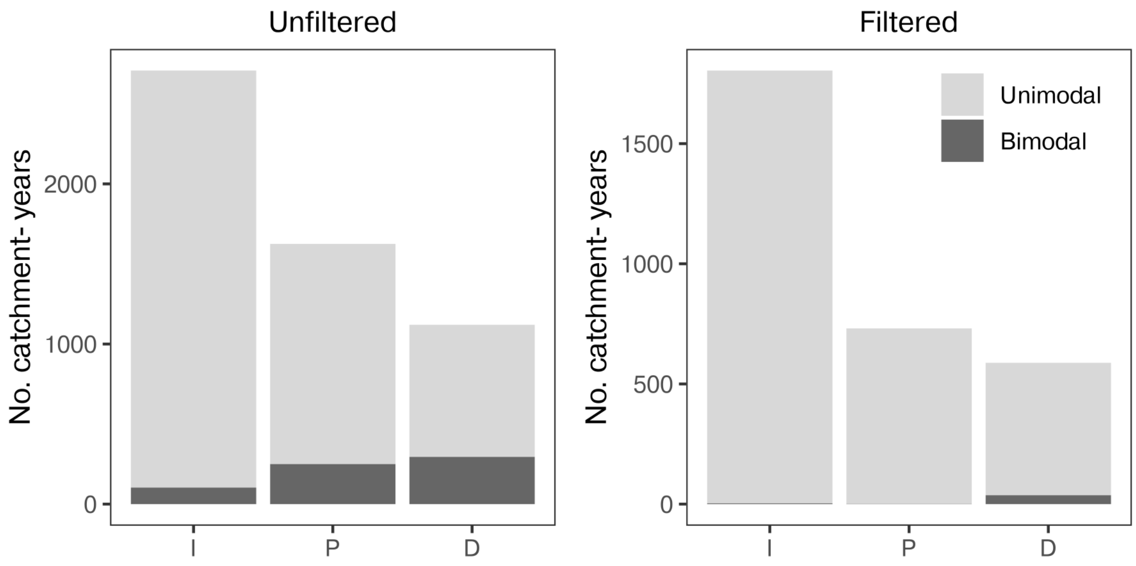


Figure S2: **Catchment-years for which unimodal and bimodal OFDs were detected.** The number of catchment-years for which significant bimodality in the OFD was detected using the Tokeshi method (Sec. 2.2) with and without filtering the data. Following filtering of demonstrably under-sampled catchment-years, the number of bimodal OFD dropped to near zero.

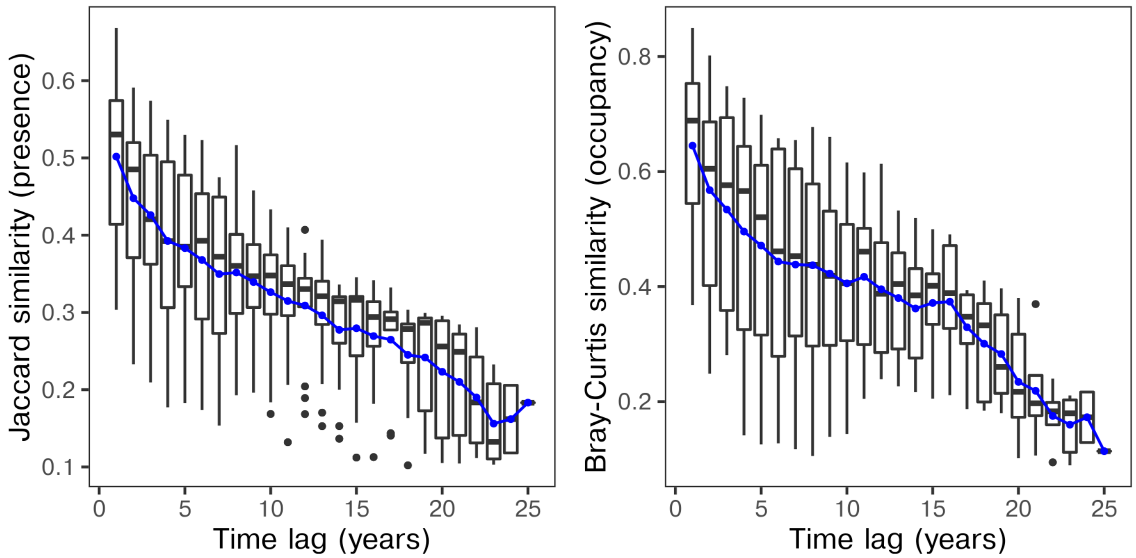


Figure S3: **The decay in similarity with temporal distance.** The decay in similarity with regards to species presence-absence (left) and site occupancy (right) at the catchment scale as a function of temporal distance (time span, in years) for a representative catchment. In Figs. 2,S6 and S7, we report the temporal decay in similarity over time inferred by averaging for a given time span (shown here in blue).

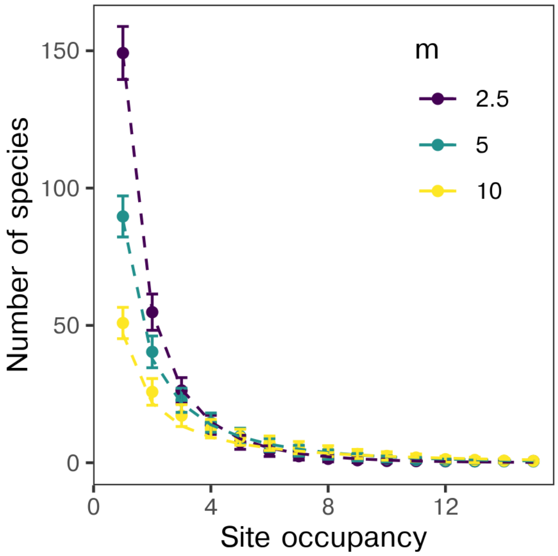


Figure S4: **Simulating the LSPOM.** The average richness in each site occupancy class across $10^3$ independent simulations of the LSPOM following $10^4$ iterations. Here $N = 50$ sites, each hosting $\alpha = 10$ local populations. Error bars indicate sample standard deviations. The mixing rate was set to $m = 2.5$, 5, and 10. Dashed lines represent least-square fits of Eq. (1) to the simulation averages. Fitted $m$ values were 2.8, 5.9 and 12.5, suggesting that the accuracy to which our method recovers $m$ from OFD is sufficient for this study.

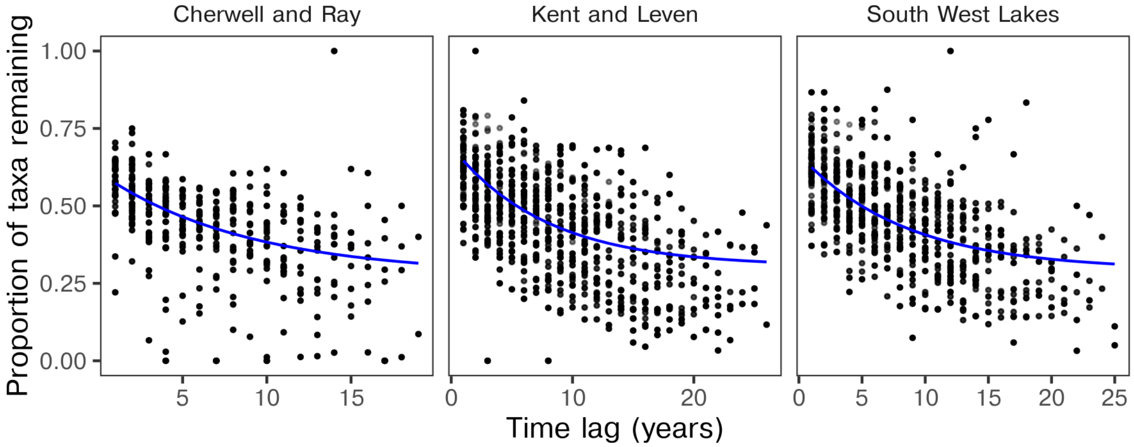


Figure S5: **Modelling catchment-scale species retention.** The detection probability and extirpation rates, assumed equal for all species in a catchment, were estimated from the parameters of an exponential decay fitted to the probability of reobserving taxa with increasing time lag. Here we show the empirical proportion of taxa remaining as a function of lag for three representative catchments from the I database and the fitted decay. Point saturation indicates weighting as described in Sec. S4.

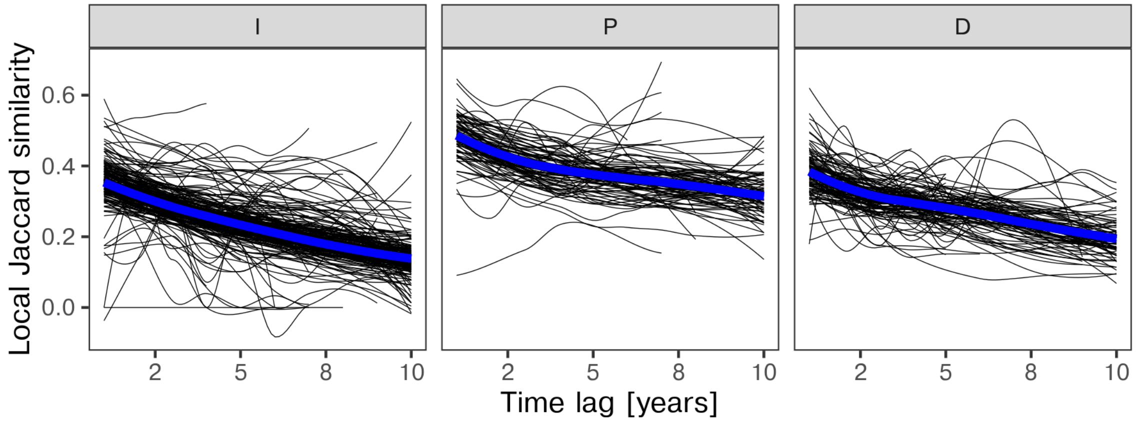


Figure S6: **Decay in Jaccard similarity by site.** The Jaccard similarity as a function of time lag for each site sampled 5 or more times. Compositional differences are considered at a maximum time lag of 10 years to account for potential changes in sampling procedure over the period. Lines represent the result of LOESS smoothing (Cleveland & Devlin, 1988) with black lines corresponding to single sites and blue lines representing the LOESS fit ignoring site identity.

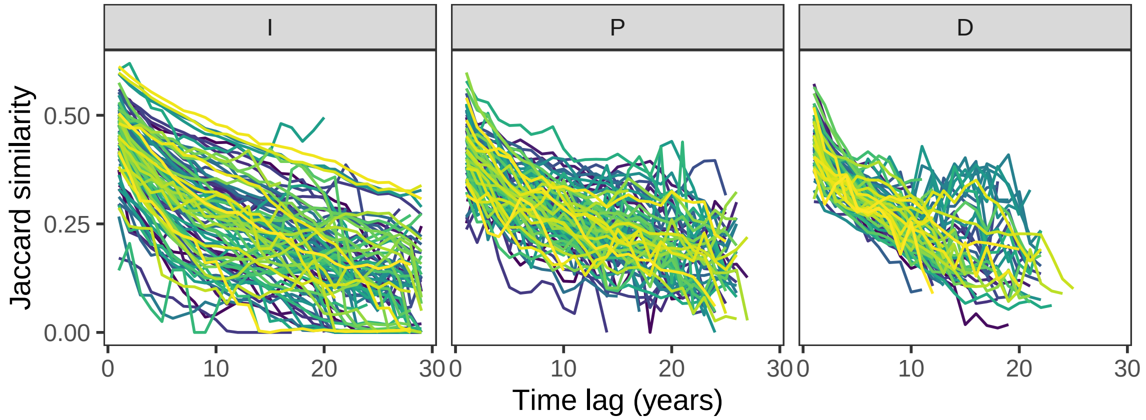


Figure S7: **Decay in Jaccard similarity by catchment.** The average Jaccard similarity as a function of time lag for each of the catchments. Jaccard similarity compares binary vectors representing presence absence. The clear negative trends shown here suggests regular colonisation and extinction at the catchment scale, though inter-annual variation in site selection may contribute to the estimated turnover rates.

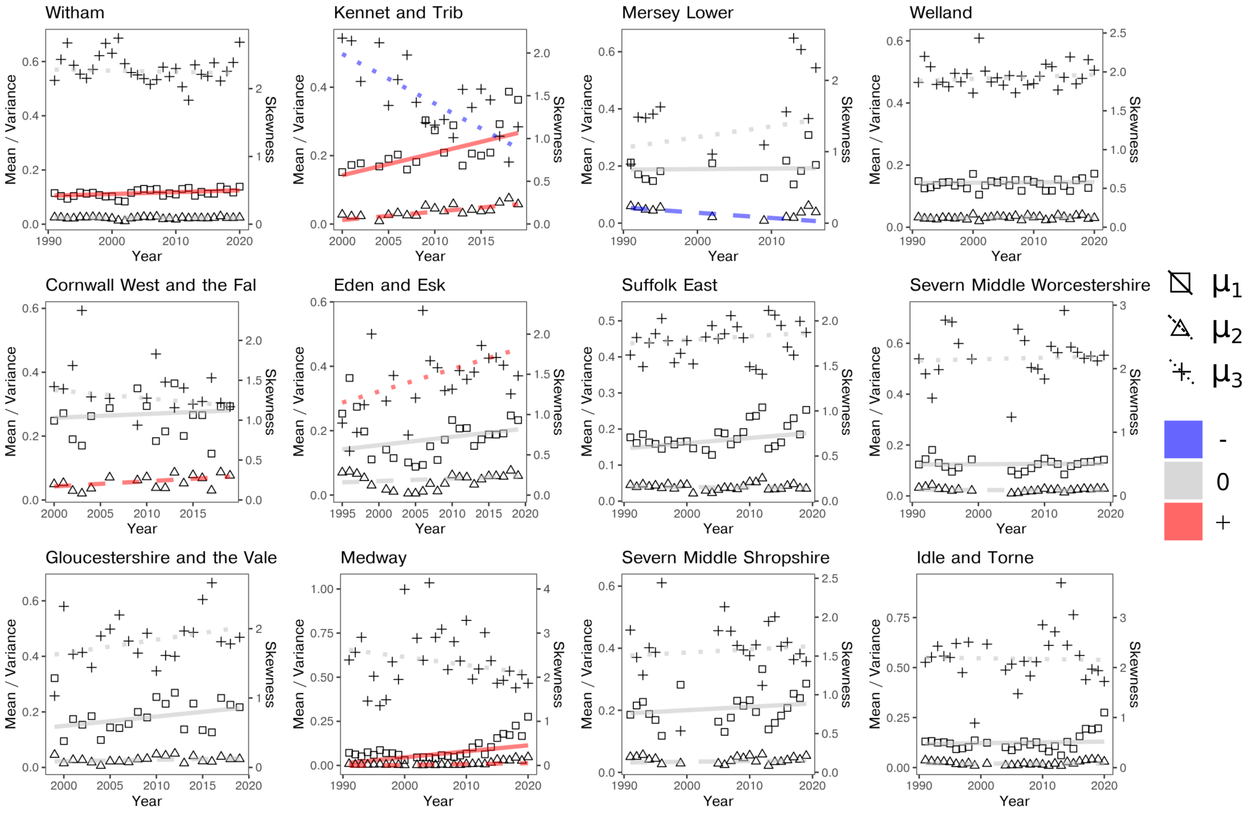


Figure S8: **Macroinvertebrate OFD moment time series for** 12 **random catchments.** Mean ($\mu_1$), variance ($\mu_2$) and skewness ($\mu_2$) of the OFD plotted against time. The slopes of the regressions are the Sen's slope estimates while the intercepts are defined as the difference between the median skewness and the median year scaled by the estimated slope (Conover, 1999). Colour indicates negative (blue), positive (red) or non-signficant slope assessed using the Mann-Kendall test of temporal trend.

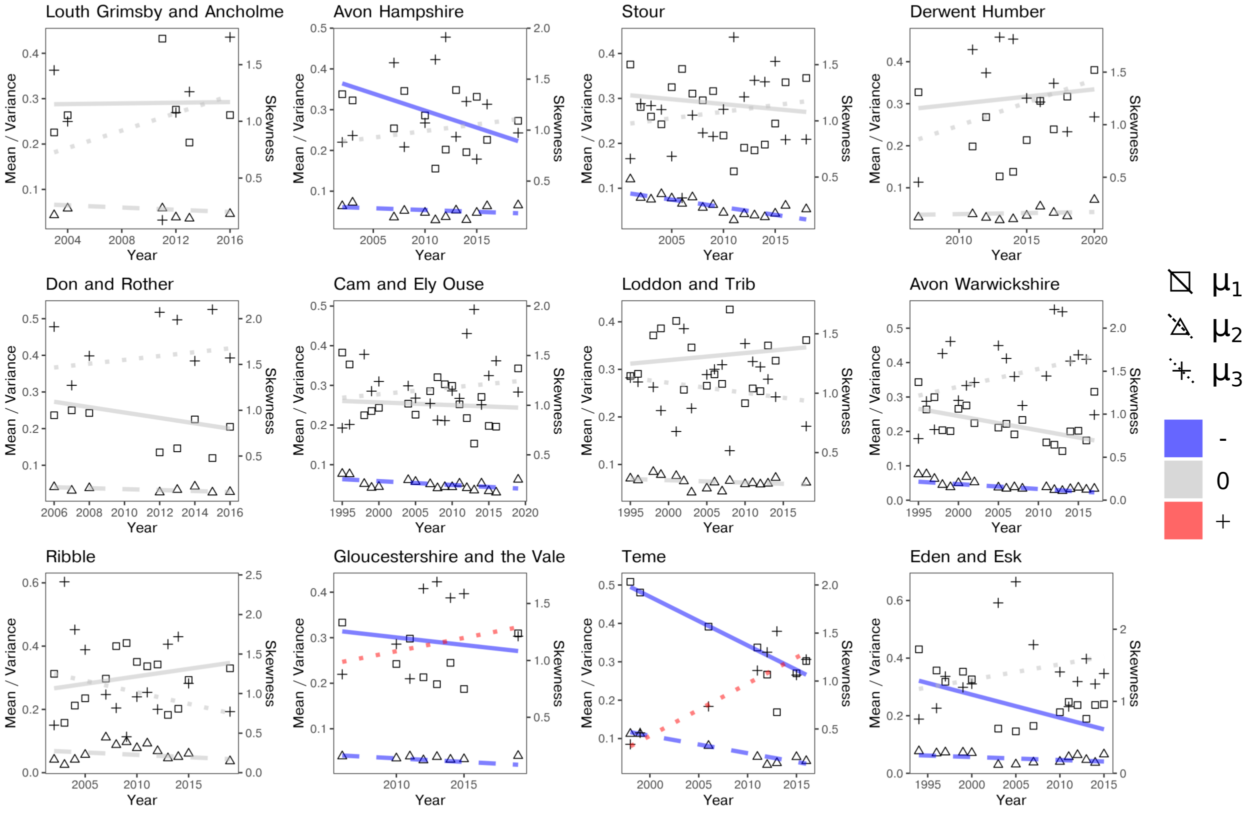


Figure S9: **Macrophyte OFD moment time series for** 12 **random catchments.** Plotted as in Fig. S8

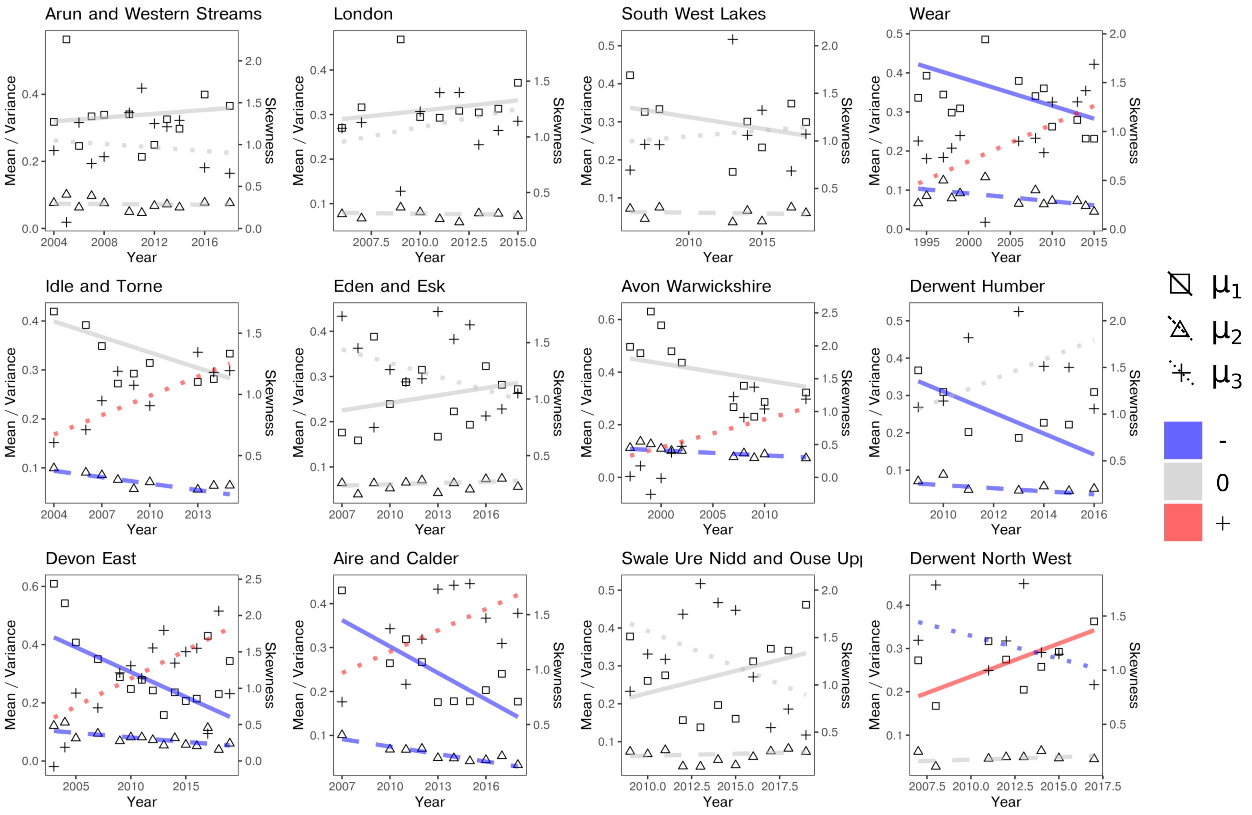


Figure S10: **Diatom OFD moment time series for** 12 **random catchments.** Plotted as in Fig. S8

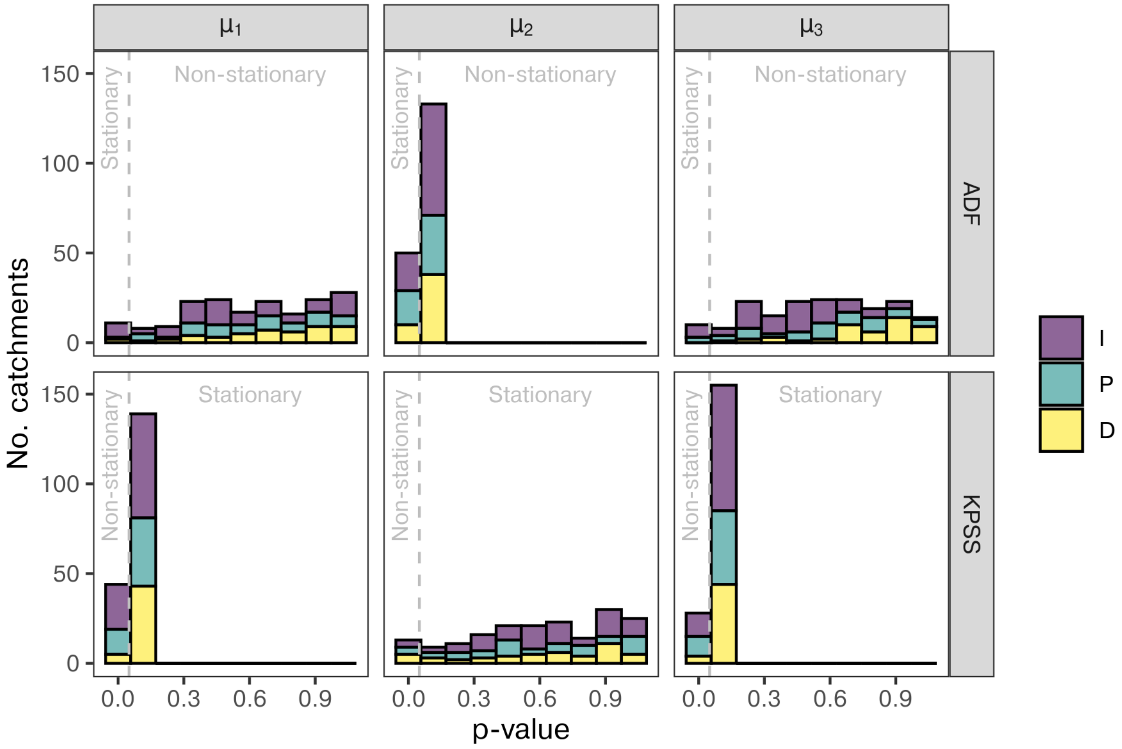


Figure S11: **The significance estimates of the ADF and KPSS tests.** The ADF and KPSS tests applied to the time series of the mean ($\mu_1$), variance ($\mu_2$) and skewness ($\mu_3$) of the catchment OFDs. Shown here are the $p$-values of the tests with $0.05$ indicated by the dashed vertical line. In the ADF test $p < 0.05$ indicates stationarity, while in the KPSS test $p > 0.05$ suggests stationarity cannot be ruled out. For the majority of the catchments $p > 0.05$ for both tests and as such the stationarity assessment was inconclusive. Interestingly, though still most often non-significant, the strongest indication of stationarity was in the case of the variance ($\mu_2$) of the OFD.

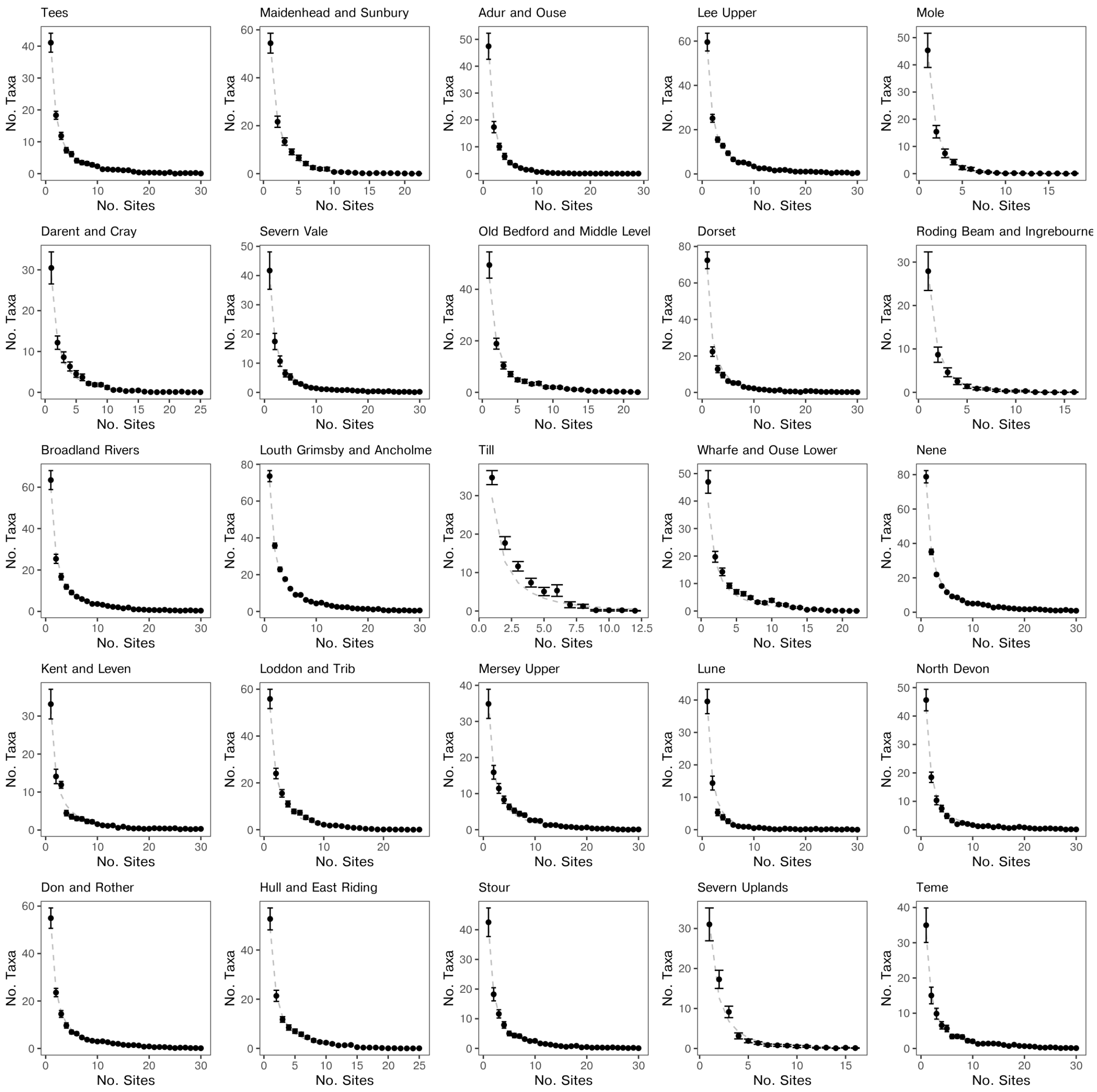


Figure S12: **Time averaged macroinvertebrate site occupancy distributions for** $25$ **random catchments.** The dashed lines represent the fitted log-series. Error bars indicate corrected standard deviation in the observed richness for a given occupancy class over the time series (without interpolation).

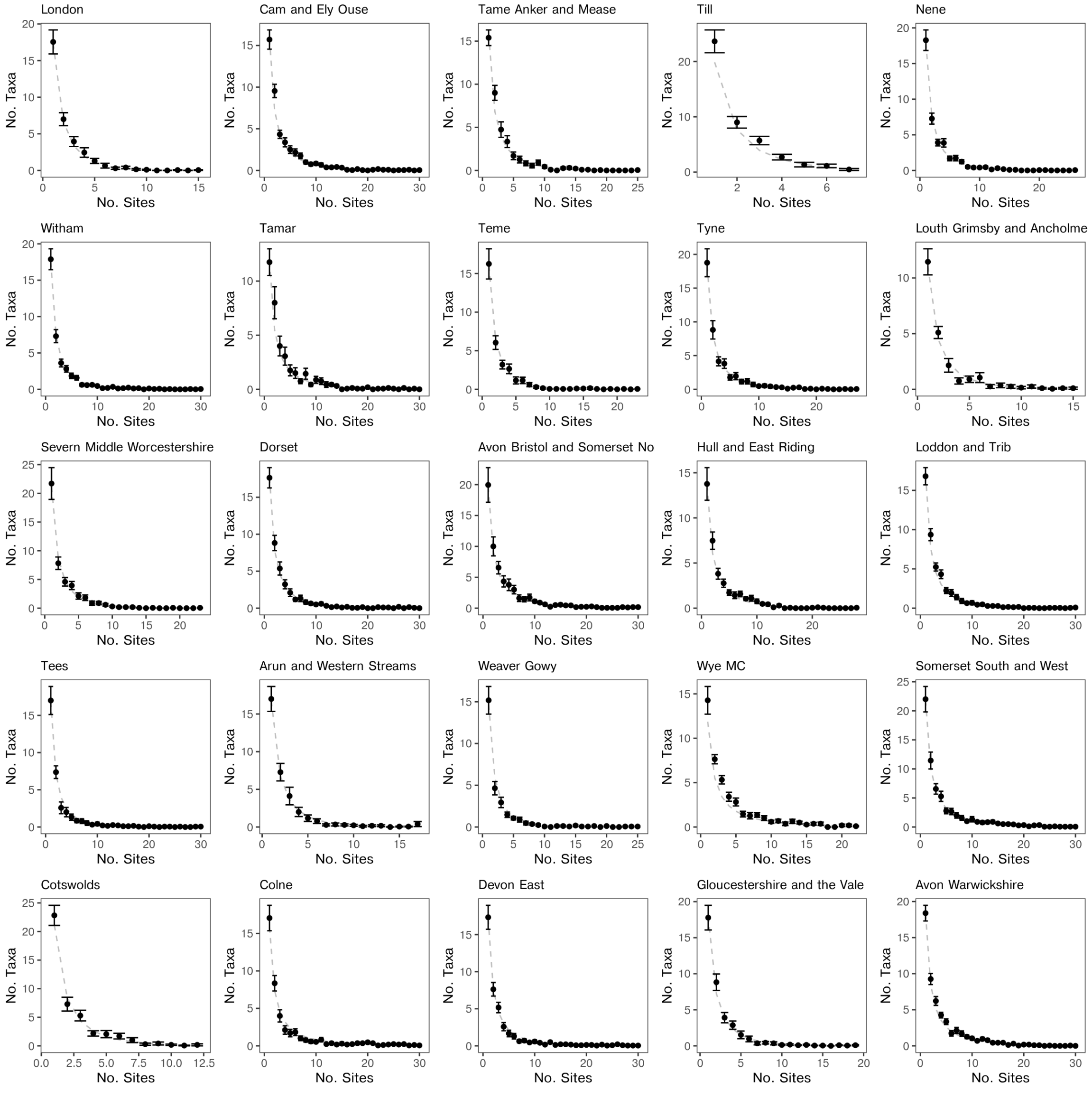


Figure S13: **Time averaged macrophyte site occupancy distributions for** $25$ **random catchments.** Plotted as in Fig. S12.

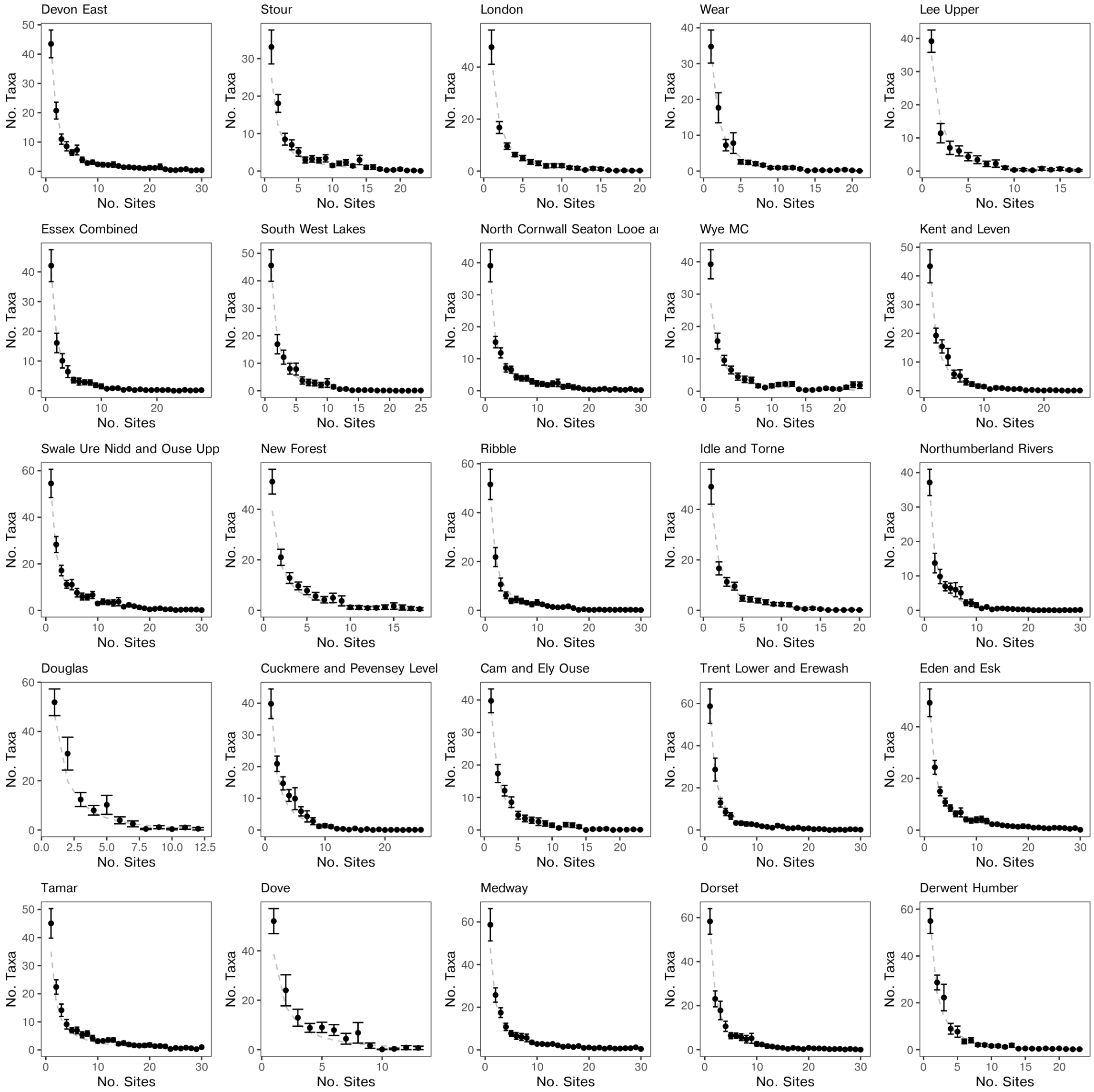


Figure S14: **Time averaged diatom site occupancy distributions for** 25 **random catchments.** Plotted as in Fig. S12.

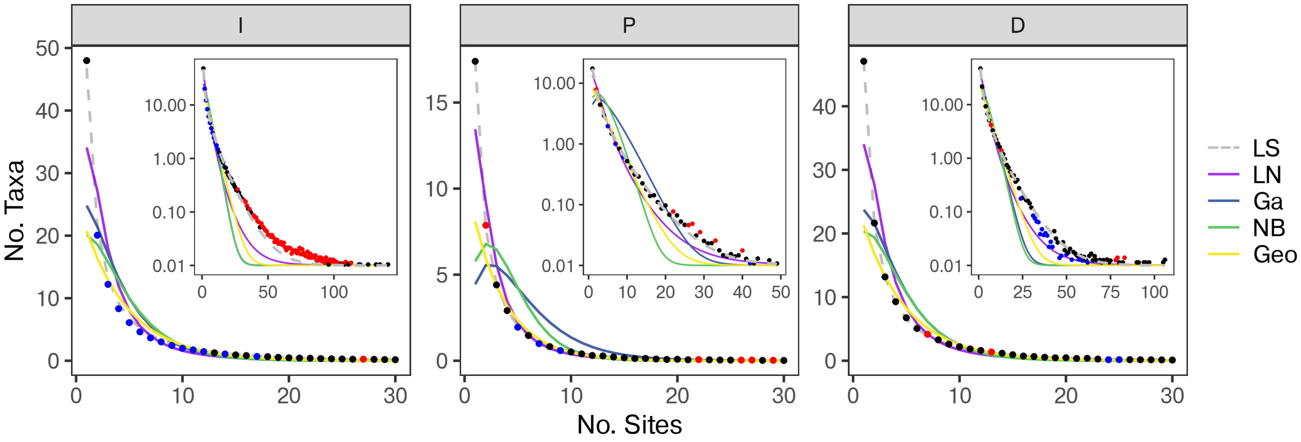


Figure S15: **Alternative models fit to national average OFDs** The catchment-time-averaged OFDs for each taxonomic group with best fitting log-series (LS), log-normal (LN), gamma (Ga), negative binomial (NB) and geometric (Geo) distributions plotted as in Fig. 2C. Here the error bars are not plotted for visual clarity. As indicated by the maximum absolute difference between average richness of each occupancy class and the various theoretical predictions (Table 2), the log-series better fits the data then any of the purely statistical functions tested.

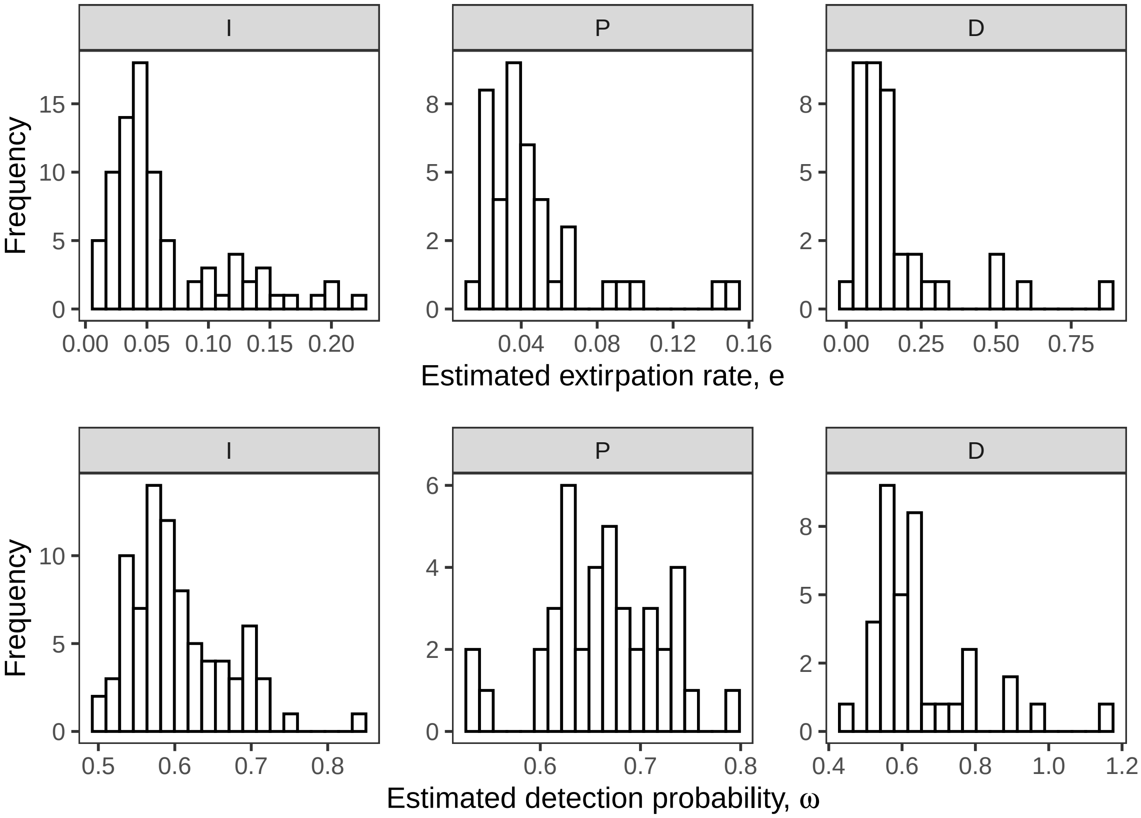


Figure S16: **Estimated extirpation and detection rates** Regional extirpation (e) and detection ($\omega$) rates estimated for each catchment as described in the text. The false negative rate is then defined as $1 - \omega$.

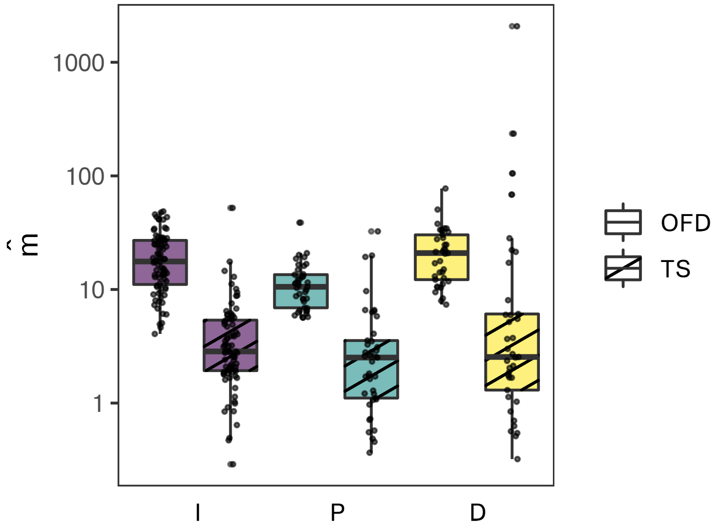


Figure S17: **Comparing mixing rates estimated from spatial and temporal analyses.** The estimated mixing rates $\hat{m}$ and the derived quantity $\hat{\alpha}$, computed by analysis of the OFD or alternatively, the observed rate of turnover (with diagonal hatching). While quantitative differences were found, these were not substantial. Further work on more systematically sampled data may reveal greater quantitative overlap between these estimates.
